# KIF1A is kinetically tuned to be a super-engaging motor under hindering loads

**DOI:** 10.1101/2022.05.16.492183

**Authors:** Serapion Pyrpassopoulos, Allison M. Gicking, Taylor M. Zaniewski, William O. Hancock, E. Michael Ostap

**Author notes:** Current address: Center for Plant Molecular Biology (ZMBP), University of Tübingen, Auf der Morgenstelle 32, 72076 Tübingen, Germany.

## Abstract

KIF1A is a highly processive vesicle transport motor in the kinesin-3 family. Mutations in KIF1A lead to neurodegenerative diseases including hereditary spastic paraplegia. We applied optical tweezers to study the ability of KIF1A to generate and sustain force against hindering loads. We used both the three-bead assay, where force is oriented parallel to the microtubule, and the traditional single-bead assay, where force is directed along the radius of the bead, resulting in a vertical force component. The average force and attachment duration of KIF1A in the three-bead assay were substantially greater than those observed in the single-bead assay. Thus, vertical forces accelerate termination of force ramps of KIF1A. Average KIF1A termination forces were slightly lower than the kinesin-1 KIF5B, and the median attachment duration of KIF1A was >10-fold shorter than KIF5B under hindering loads. KIF1A rapidly reengages with microtubules after detachment, as observed previously. Strikingly, quantification enabled by the three-bead assay shows that reengagement largely occurs within 2 ms of detachment, indicating that KIF1A has a nearly tenfold faster reengagement rate than KIF5B. We found that rapid microtubule reengagement is not due to KIF1A’s positively charged loop-12; however, removal of charge from this loop diminished the unloaded run length at near physiological ionic strength. Both loop-12 and the microtubule nucleotide state have modulatory effects on reengagement under load, suggesting a role for the microtubule lattice in KIF1A reengagement. Our results reveal adaptations of KIF1A that lead to a novel model of super-engaging transport under load.

## Introduction

KIF1A is a cytoskeletal motor in the kinesin-3 family that transports intracellular cargo in axons and dendrites (1, 2). A number of human mutations in KIF1A have been identified that lead to neurodegenerative diseases, termed KIF1A Associated Neurological Disorders (KAND) (3, 4). KIF1A is functionally distinctive in the kinesin superfamily in that it has a fast-stepping rate and enhanced processivity in the absence of mechanical loads compared to other characterized motors. A positively charged loop-12 insert, the “K-loop,” is unique to the kinesin-3 family and has been linked to the motor’s superprocessive behavior (Fig. 1A) (5, 6). However, single-molecule experiments found the K-loop did not contribute to the superprocessivity of KIF1A dimers at low ionic strength (7–9). Despite being superprocessive, mechanical loads that resist plus end-directed stepping cause KIF1A to detach from the microtubule more readily than the well-studied kinesin-1 (10–13).

**Figure 1:**
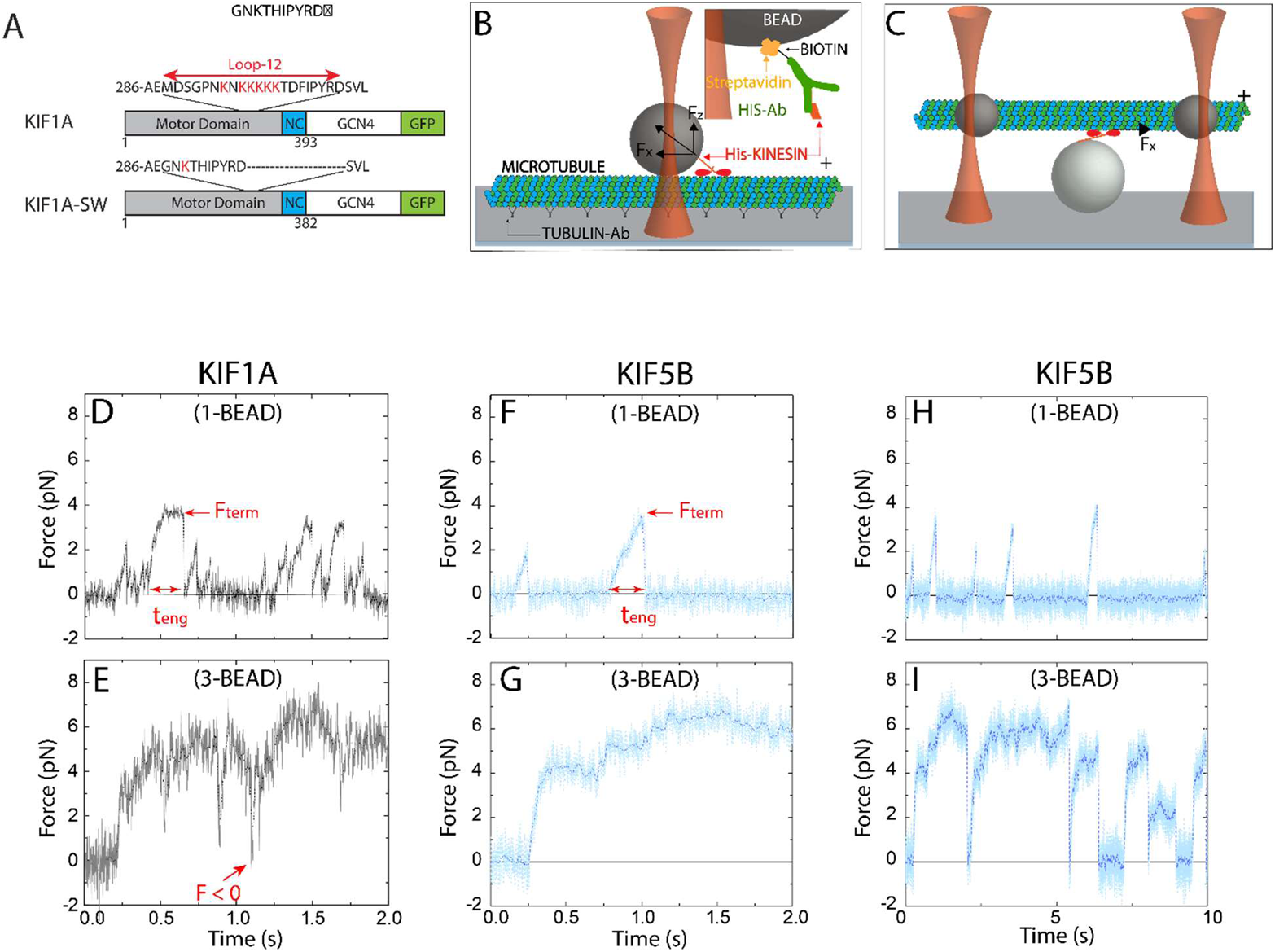
KIF1A performance in single-bead and three-bead assays. (A) KIF1A constructs used in this study. Wild-type KIF1A consists of the motor domain the neck coil region (NC) of rat KIF1A, followed by a leucine zipper dimerization domain (GCN4) and a C-terminal GFP and His_6_-tag. In the KIF1A-SW swap mutant, the loop-12 was replaced by the corresponding sequence of *Drosophila* kinesin-1. (B and C): Diagram of the (B) single-bead and (C) three-bead assays with attachment strategy (inset); beads are not drawn to scale. (D and E): Representative force traces of KIF1A at 2 mM ATP in (D) the single-bead assay and (E) the three-bead assay. F_term_ is defined as the force at the termination of a force ramp and t_eng_ is defined as the duration of a force ramp. Red arrow in (E) highlights instance of force after termination of a force ramp crossing the zero-force baseline before initiating a new force ramp. (F and G): Force traces of KIF5B using (F) the single-bead assay and (G) the three-bead assay with the same time axis as D and E. (H and I): KIF5B force traces at expanded time scale, showing larger intervals between force ramps and long force plateaus in the three-bead geometry.

A recent biochemical study exploring the mechanochemical adaptations of KIF1A suggested that rear-head detachment is an order of magnitude faster than found for kinesins-1 or -2, and that this feature helps to explain its rapid stepping rate. This kinetic feature also results in a predominant steady-state intermediate that is bound via a single “weakly-bound” post-hydrolysis motor domain through electrostatic interactions with the microtubule (14). This single-head microtubule interaction may result in a molecule that is vulnerable to detachment under mechanical load. Indeed, a recent study using a single-bead optical tweezer found that KIF1A bound for relatively short durations under load and generated stall forces of 3.1 pN, compared to 4.6 pN for kinesin-1 (13). Interestingly, KIF1A was found to recover processive stepping after detachment more readily than kinesin-1, and this property has been attributed to the unique K-loop (8, 14).

The specific sequences and biochemical tuning that underlie the superprocessivity and force sensitivity of KIF1A are still under investigation. The α4-helix, which forms a substantial part of the microtubule binding interface, is conserved between the kinesin-1 and kinesin-3 families, but there are positively charged residues in loop-8, loop-11, and the α6-helix of KIF1A that, when substituted for their kinesin-1 counterparts, reduce the unloaded run length substantially (15). Furthermore, the N-terminal cover strand of kinesin-3, which stabilizes the docked neck linker and contributes to force generation, is shorter than that of kinesin-1 and forms a less extensive hydrogen bonding network with the motor domain and neck linker (13, 16). These unique structural features within the catalytic core and at the motor-microtubule interface raise the possibility that KIF1A motor kinetics are affected by force differently than kinesin-1.

Given the unique connection between the neck-linker and motor of KIF1A, it is important to consider how the geometry of forces applied to the motor affects its mechanochemistry. The single-bead assay (Fig. 1B), which is commonly used for measuring the force generated by kinesin motors (including KIF1A (13, 17)), introduces a vertical component to the force applied to the kinesin due to contact of the bead with the underlying surface-immobilized microtubule (18, 19). This vertical force component acts to separate the motor from the microtubule. Vertical forces can be minimized by using a three-bead assay (Fig. 1C), in which the motor is attached to a surface-immobilized bead and a microtubule “dumbbell” is held above it by two laser-trapped beads attached near the microtubule ends (19, 20). In recent work, it was found that the microtubule detachment rate of human kinesin-1, KIF5B, was substantially slower in the three-bead assay (19), suggesting that the vertical force inherent to the single-bead assay contributes to the measured motor detachment kinetics. Thus, given the recent finding that KIF1A detaches from microtubules more readily under force (13), it is important to examine the contribution of parallel and vertical forces to processive stepping.

In the present work, we investigated the performance of KIF1A in single- and three-bead optical trap assays and compared its performance to kinesin-1. We found that although KIF1A can achieve forces up to 6 pN, it is not a superprocessive motor under load. Rather, it is super-engaging, in that under opposing forces it readily disengages from the microtubule, but it quickly reengages and initiates a new force ramp within 2 ms. We also found that at near physiological ionic strengths, the K-loop contributes substantially to the unloaded run length, but only minimally to the load-dependent detachment kinetics. These results suggest that during vesicle transport in cells, where forces are predominantly oriented parallel to the microtubule, KIF1A is able to detach and rapidly recover motility under load, an adaptation that facilitates bidirectional transport and navigation around obstacles.

## Results

### KIF1A generates comparable forces to KIF5B and reengages more frequently with the microtubule

To probe the force generating properties of KIF1A, we used optical tweezers in both a single-bead and three-bead configuration (Fig. 1B and C) at saturating ATP (2 mM). Because full length KIF1A molecules adopt an auto-inhibited conformation (3, 21, 22), we used a *Rattus norvegicus* KIF1A construct consisting of the motor and neck coil domains dimerized by a GCN4 leucine zipper and followed by a GFP (7) (Fig. 1A). KIF1A concentrations used in the optical tweezers experiments were sufficiently low to ensure that observed interactions are due to single KIF1A dimers (see Materials and Methods).

In the single-bead assay, KIF1A molecules pulled the bead out of the center of the stationary optical trap to forces of ~ 4 pN (Fig. 1D). Terminations of force ramps were followed by strictly monotonic decreases in force as the bead relaxed back toward the center of the optical trap. By averaging many such events, we found that the relaxation time was ≤ 2 ms, which is near the expected relaxation time of a single bead in the absence of any interactions with the microtubule (see Supporting Information; Fig. S1) (23). These rearward displacements may reflect complete dissociation of KIF1A from the microtubule; alternatively, they could reflect KIF1A slipping backwards while maintaining weak association with the microtubule, as shown previously for KIF5B (24, 25). As we cannot differentiate between these attachment states, we refer to the force value at the termination of each force ramp as the *termination force* (F_term_; Fig. 1D and F). We refer to this transition at F_term_ as *disengagement* of the motor from the microtubule.

Following disengagement, KIF1A quickly reengages with the surface-immobilized microtubule and resumes forward motion (Fig. 1 D). Successive KIF1A force ramps were more closely spaced in time than those measured for KIF5B under identical assay conditions. The maximal KIF1A termination forces (F_term_) of ~4 pN were lower than KIF5B, and the duration of the force ramps, defined as the engagement time (t_eng_), were shorter for KIF1A than for KIF5B (Fig. 1D and F). The lower forces and rapid reengagement kinetics of KIF1A agree with a recent single-bead optical trapping study using a rat KIF1A construct (13).

In the single-bead assay, forces are applied to kinesin in directions both parallel and normal to the long-axis of the microtubule (Fig. 1B; (18, 19)). To investigate the force generating properties of KIF1A in the absence of this normal force component, we used the three-bead assay, in which the motor is attached to a surface-immobilized bead and a microtubule “dumbbell” is held above it by two laser-trapped beads attached near the microtubule ends (Fig. 1C; Materials and Methods). In the three-bead assay KIF1A developed maximal forces of ~6 pN, substantially larger than in the single-bead assay and close to the forces generated by KIF5B (see Results below). Notably, the durations of the KIF1A force ramps were still substantially shorter than observed for KIF5B (Fig. 1E and G). After disengaging, KIF1A rapidly reengaged and initiated the next force ramp before the dumbbell fully relaxed to the zero-force baseline. This rapid reengagement was rarely observed for KIF5B (Fig. 1G and I).

### In the absence of vertical forces KIF1A generates large pulling forces and repetitively reengages with the microtubule

To isolate the influences of vertical and horizontal forces on KIF1A stepping, we quantified the force generating capacity of KIF1A and the microtubule reengagement kinetics following termination of a force ramp. A representative example of a long (>100 s) trace that contains many consecutive KIF1A force ramps is shown in Fig. 2A. When the distribution of instantaneous forces was plotted (Fig. 2B), two clear modes were apparent: a peak around the zero-force baseline, and a peak around the average force where force ramps terminated. For our analysis, force ramps that initiated at forces within two standard deviations of the zero-force baseline were termed *primary events*, whereas force ramps that initiated at forces greater than two standard deviations from the baseline were termed *secondary events* (Fig. 2C and D). For KIF1A, 39% of the force ramps in the three-bead assay qualified as secondary events, whereas only 11% qualified as secondary events in the single-bead assay (Table S1). This difference may result from the microtubule remaining near the immobilized motor in the three-bead trap, whereas the motor position is less constrained in the single-bead trap due to potential rotation of the bead. It is also possible that tensile forces applied by the two traps on the microtubule in the three-bead assay (see Materials and Methods) may deform the microtubule lattice and thereby enhance motor re-engagement kinetics (26).

**Figure 2:**
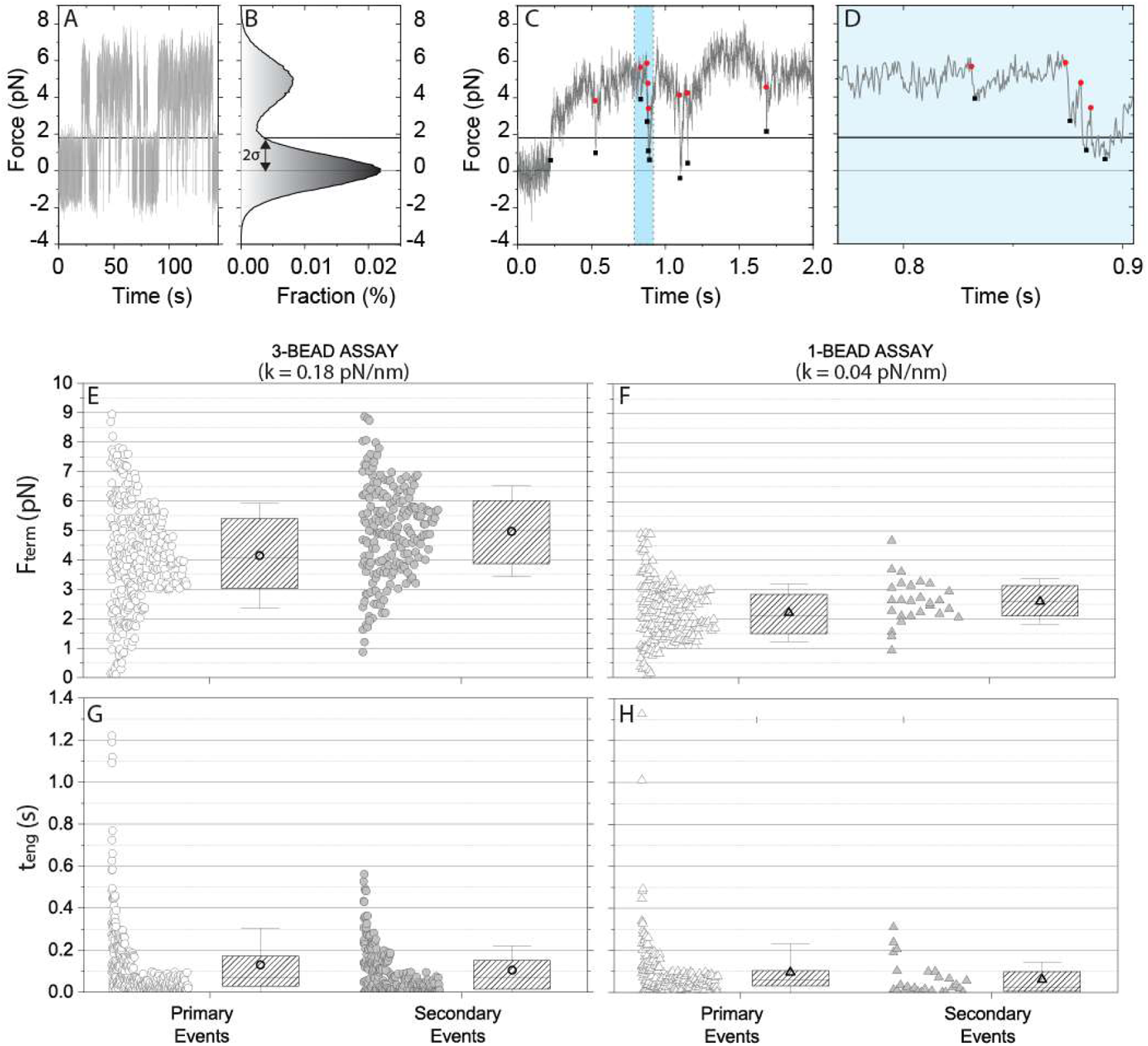
Quantification of KIF1A primary and secondary force ramps. (A) Long duration force trace of KIF1A in the three-bead assay. (B) Corresponding stationary distribution of force, exhibiting a peak at the zero-force equilibrium position and a ramp force peak at 5 pN. Primary events are defined as force ramps that start within two standard deviations of the zero-force baseline (horizontal 2σ line at F = 1.8 pN). (C) Sample force trace in the three-bead assay showing multiple disengagement events (red dots) and reengagement events (black squares). Reengagements that initiate below the 2σ line are considered primary binding events, whereas events that start above the threshold are considered secondary events. (D) Expanded view of highlighted portion of (C). (E and F): KIF1A termination forces, F_term_ for primary and secondary events in the (E) three-bead and (F) single-bead assays. In each case, raw data are shown at left and the average (open circle and triangle), median (horizontal line), quartile (boxes) and standard deviation (error bars) shown at right. (G and H): Engagement durations, t_eng_ for primary and secondary events in the (G) single-bead and (H) three-bead assay. Raw data are shown at left and the mean (open circles and triangles), quartile (shaded box) and standard deviation (error bars) are shown at right.

**Table 1:**
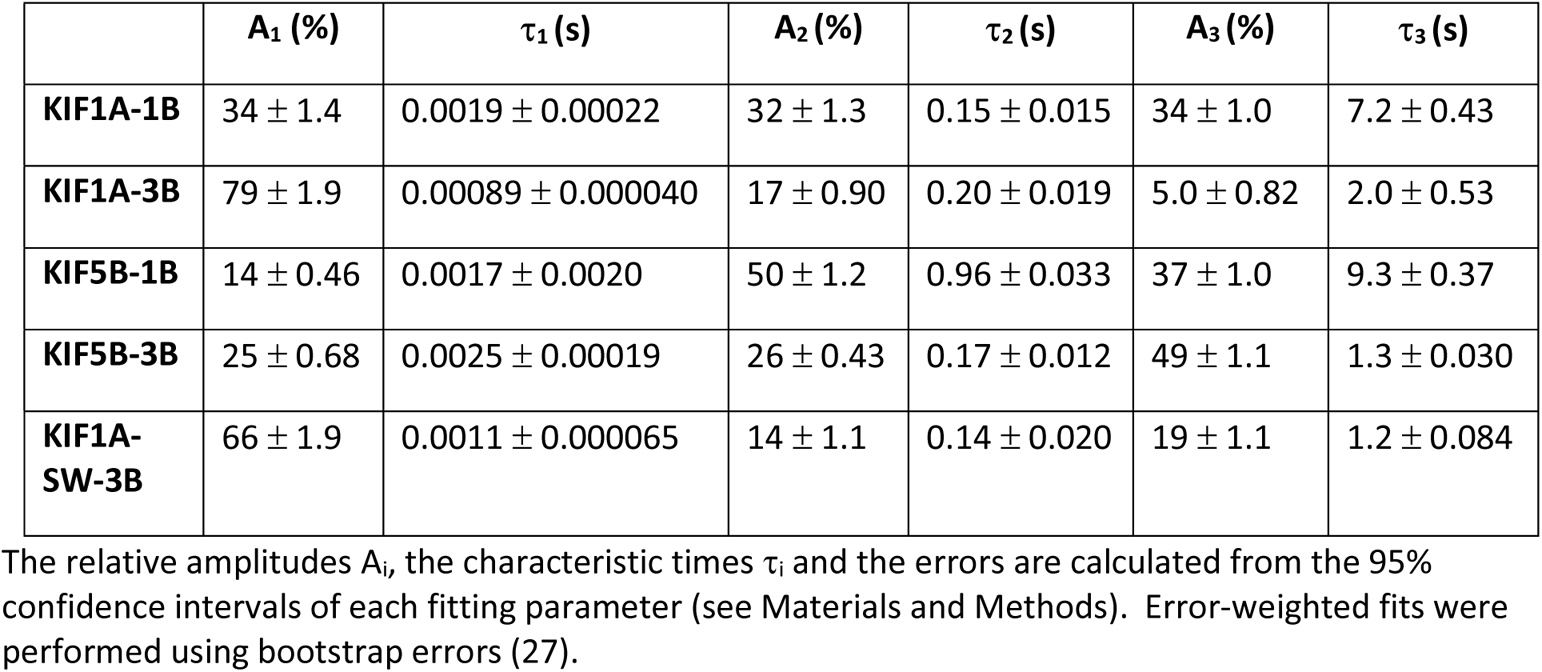
Fitting results of cumulative probability of reengagement using a tri-exponential function

To characterize the mechanical performance of KIF1A against hindering loads, we quantified the distributions of termination forces, F_term_, and the durations that motors engaged with microtubules before termination of a force ramp, t_eng_. For both the single-bead and three-bead assay, <F_term_> was slightly higher for the secondary events relative to the primary ones (Fig 2E and F; Table S2). The higher forces may be expected, because primary events begin at lower initial forces, and hence require more time to build to higher forces; it also reflects the fact that motors usually disengaged before reaching a stable force. To better quantify disengagement kinetics, we compared the distribution of motor engagement times, t_eng_, in the three-bead assay (Fig. 2G and H; values in Table S2). The median t_eng_ was similar for primary and secondary events, at 69 ms and 67 ms, respectively, consistent with primary and secondary events reflecting similar motor engagement processes and differing only in their initial starting positions. In the single-bead assay, the primary events had similar median durations as the three bead (62 ms), whereas the secondary events were fairly rare and were shorter duration (23 ms).

Because the microtubule dumbbell was pre-tensioned to reduce thermal noise, higher trap stiffnesses were used with the three-bead assay. The higher stiffness also compensated for the larger viscous drag of the dumbbell compared to a single bead, resulting in the relaxation times being similar for the two assays (see Supplemental Information and Fig. S1). As a result of this higher stiffness, the loading rate during force ramps (dF_x_/dt = k_x_·v_x_) was faster in the three-bead assay, raising the possibility that the lower termination forces in the single-bead assay may result from the longer time required to generate high forces. To rule out this possibility, we repeated the single-bead assay at a higher trap stiffness and found that, although F_term_ increased slightly, it was still substantially lower than the value for the three-bead assay (Fig. S2A). Furthermore, using a more comparable trap stiffness in the single-bead assay, the median engagement time fell to 35 ms, highlighting shorter engagement times in the single-bead assay (Fig. S2B). Therefore, termination forces were higher in the three-bead assay than in the single-bead assay, consistent with the vertical forces inherent in the single-bead assay limiting the duration of the force ramps.

### KIF1A engagement times are short under load and reengagement occurs within milliseconds

A consistent feature of KIF1A behavior in both the single- and three-bead assays was the rapid reengagement of the motor with the microtubule following the termination of a force ramp. This behavior was observed previously in a single-bead study (13), but not quantified. To characterize this reengagement behavior, we determined the restart time, t_restart_, defined as the time between termination of one force ramp and initiation of the next. For KIF1A and KIF5B in the two optical trapping geometries, the cumulative probability distribution of restart times showed a population of fast restart events on the ms timescale and two slower populations with time constants > 100 ms (Fig. 3A and B). The cumulative distributions of t_restart_ were fitted to the sum of three exponentially distributed populations. In the three-bead assay, the time constant of the fastest phase was 0.89 ms for KIF1A and 2.5 ms for KIF5B (Table 1), which are on the order of the dead time of the experiment set by the relaxation time of the trapped beads (see Methods). Strikingly, 79% of KIF1A reengagement events occur within the fast phase, compared with only 25% of KIF5B reengagements (Table 1). For both motors, the amplitude of the fast phase is smaller for the single bead assay (Table 1), suggesting that the assay geometry significantly impacts the reengagement times (Fig. 3B). The slower phases are likely due to motors detaching from the microtubule with reengagement being limited by the steric constraints of the experimental geometry and the motor kinetics.

**Figure 3:**
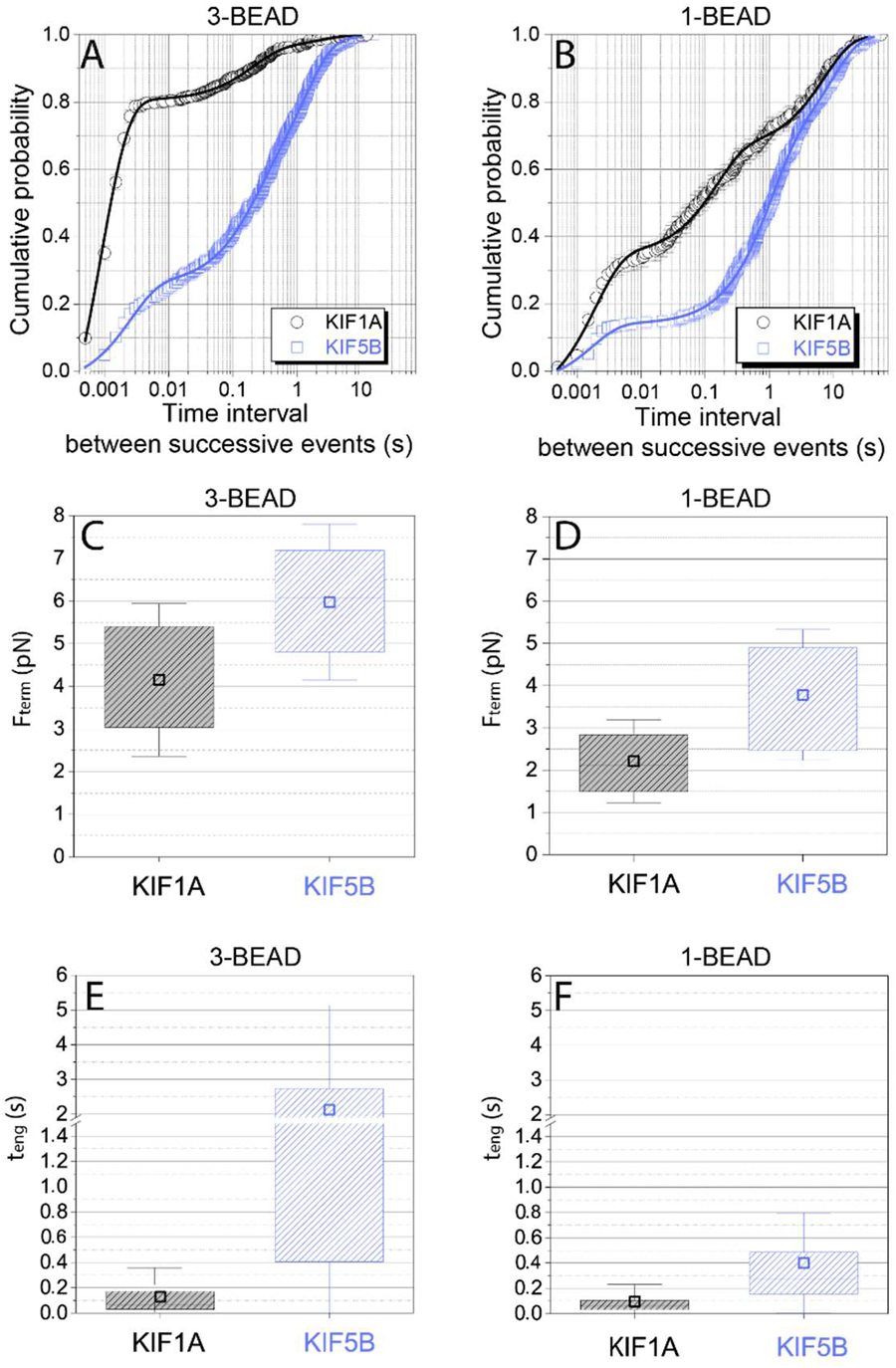
Comparison of KIF1A and KIF5B engagement/disengagement dynamics. (A and B): Cumulative probability distribution of time intervals between successive force ramps (t_restart_) for KIF1A and KIF5B in the (A) three-bead and (B) single-bead assays. Data include both primary and secondary events. Error bars are calculated using the bootstrap method (27), and solid lines represent fitting to a three-exponential function with offset (see Materials and Methods). (C and D): Comparison of KIF1A and KIF5B termination forces, F_term_ for primary events in the (C) three-bead and (D) single-bead assays. Data are presented as mean (open squares), median (horizontal line), quartiles (shaded boxes), and standard deviation (error bars). (E and F): Comparison of KIF1A and KIF5B engagement durations, t_eng_ for primary events in the (E) three-bead and (F) single-bead assays. Note the break introduced at 1.5 s in the y-axis due the large difference between the median values of t_eng_ between KIF1A (0.069 s) and KIF5B (1.1 s) in the three-bead assay.

To compare the ability of the two motors to generate and sustain forces against hindering loads oriented exclusively parallel to the microtubule, we compared the average F_term_ and median t_eng_ in the three-bead assay. As shown in Fig, 3C, <F_term_> was ~6 pN for KIF5B, but was only ~4 pN for KIF1A. This reduced capacity of KIF1A to generate and sustain forces was also seen in the single-bead assay (Fig. 3D). Therefore, even though KIF5B steps at less than half the speed of KIF1A and thus takes longer time to generate large forces, the KIF5B force ramps terminate at higher forces. Consistent with these higher forces, the median engagement time during force ramps in the three-bead assay was more than an order of magnitude shorter for KIF1A than for KIF5B (Fig. 3E), and shorter engagement times for KIF1A were also seen in the single-bead assay (Fig. 3F). In summary, KIF1A disengages from the microtubule under load more readily than KIF5B, and consequently only ~15% of KIF1A ramps reach 6 pN, whereas ~50% of KIF5B ramps reach and exceed 6 pN (Fig. S3).

### The KIF1A loop-12 contributes to superprocessivity but does not enhance initial landing on microtubules

A distinctive feature of the KIF1A motor domain is a loop-12 insert containing six positively charged lysines that are thought to interact electrostatically with the negatively charged C-terminal tail of tubulin (5, 6) (Fig. 1A). To test the contribution of loop-12 to the microtubule engagement duration and superprocessivity of KIF1A at near physiological ionic strength, we made a loop swap mutant, KIF1A-SW, by exchanging the native KIF1A loop-12 that contains the six lysines for loop-12 from *Drosophila* kinesin-1, which contains only one lysine (Fig. 1A). Single-molecule TIRF experiments in 80 mM PIPES buffer showed that in the absence of external forces, KIF1A and KIF1A-SW move along surface immobilized microtubules at similar average speeds <V> of 1.2 ± 0.36 μm/s and 1.3 ± 0.42 μm/s, respectively (Fig. 4A-C; Table S3). Notably, the average run length <RL> of KIF1A-SW (1.1 ± 0.56 μm) was approximately six-fold lower than for KIF1A (6.3 ± 4.2 μm) (Fig. 4D; Table S3).

**Figure 4:**
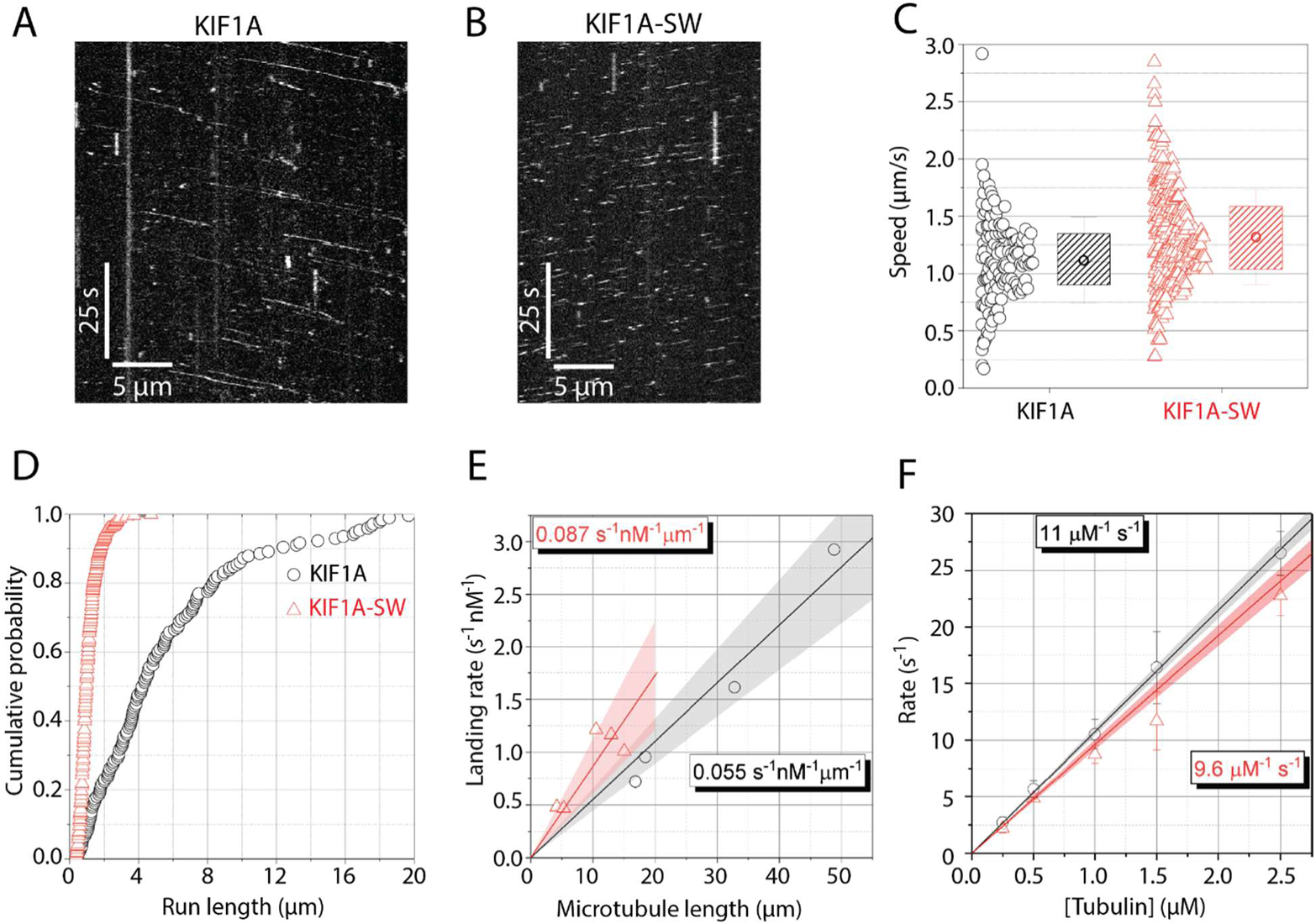
Influence of loop-12 on KIF1A performance under zero load. (A and B) Kymographs from single-molecule TIRF assays of (A) KIF1A and (B) KIF1A-SW on control microtubules in 2 mM ATP. (C) Comparison of single-molecule speeds between KIF1A (black circles) and KIF1A-SW (red triangles), showing raw data at left and mean (open symbols), median (horizontal line), quartile (shaded box) and standard deviation (error bar) at right. (D) Cumulative probability distribution of KIF1A and KIF1A-SW run lengths, using the same symbols and colors as in (C). (E) Plot of the single-molecule landing rate (s^−1^·nM^−1^) for KIF1A (black circles) and KIF1A-SW (red triangles) as a function of the microtubule length (μm). The solid lines are linear fits, which correspond to the landing rates (s^−1^·nM^−1^·μm^−1^) for each motor. Shaded areas represent the 95% confidence bands of the linear fits. (F) Plot of the mantADP release rate upon mixing mantADP bound motors with different concentrations of taxol-stabilized microtubules by stopped-flow. The linear fits to the data represent the bimolecular on-rate for KIF1A binding to microtubules (μM^−1^·s^−1^). Shaded areas represent the 95% confidence bands of the linear fits.

Previous studies that used a loop-swap construct found that swapping the lysine containing region of loop-12 of KIF1A had minimal effect on the run length at low ionic strength (12 mM PIPES buffer), although it did enhance the landing rate of KIF1A (8). To resolve this discrepancy, we carried out single-molecule experiments in 12 mM PIPES buffer and found that the run lengths of KIF1A and KIF1A-SW were similar to one another, consistent with the previous studies (Fig. S6). Thus, we conclude that at near-physiological ionic strength, positive charge in loop-12 contributes to the KIF1A run length, but at low ionic strength the loop swap has a negligible effect on the run length.

In contrast to the different run lengths, we found that KIF1A and KIF1A-SW had similar single-molecule microtubule landing rates in 80 mM PIPES, which were 0.055 ± 0.011 s^−1^·μM^−1^·μm^−1^ and 0.087 ± 0.026 s^−1^·μM^−1^·μm^−1^, respectively (mean ± 95% confidence interval; Fig. 4E). Because this single-molecule landing rate method is highly sensitive to differences in relative activity between different motor preps, we performed complementary stopped-flow experiments to determine the apparent second-order rate constant for microtubule binding. In this assay, KIF1A motors preincubated with mantADP are rapidly mixed with a range of microtubule concentrations in the presence 1 mM ATP. Microtubule binding to KIF1A in the presence of excess ATP results in irreversible mantADP release, resulting in a decrease in mant fluorescence. Consistent with the single-molecule landing rates in 80 mM PIPES, the bimolecular on-rates of KIF1A (10.6 ± 0.5 μM^−1^s^−1^) and KIF1A-SW (9.6 ± 1.5 μM^−1^s^−1^) were similar (Fig. 4F).

To further investigate the interaction of the KIF1A loop-12 with microtubules, we removed the negatively charged C-terminal tail of tubulin by subtilisin proteolysis (Materials and Methods; Fig. S4). We found that the average speed of KIF1A on subtilisin microtubules was unaffected (1.3 ± 0.39 μm/s; Table S3), but the average run length was decreased by ~5-fold to 1.3 μm (494043v1; Table S3). Thus, decreasing the charge of the KIF1A loop-12 or cleaving the C-terminal tubulin tail had similar effects, implicating electrostatic interactions between these regions as an important contributor to the superprocessivity of KIF1A under zero load. In summary, at the near-physiological ionic strength used in this study, the highly-charged loop-12 is necessary for the unloaded superprocessivity of KIF1A, but it is not required for the initial strong-binding of KIF1A to microtubules.

### Both the loop-12 and the nucleotide state of the microtubule affect the load-dependent properties of KIF1A

Given the importance of loop-12 for the unloaded processivity of KIF1A at near-physiological ionic strength, we investigated the motile properties of KIF1A-SW under load. In the three-bead assay the median motor engagement duration, median-t_eng_, decreased from 0.069 s for KIF1A to 0.039 s for KIF1A-SW (Man Whitney test p < 0.001). Consistent with these shorter engagement times, the mean termination force, <F_term_>, decreased from 4.1 pN for KIF1A to 3.5 pN for KIF1A-SW (Fig. 5; Table II). To determine if these lower termination forces were caused by differences in the motor stepping rate under load, force-velocity profiles were compared for KIF1A and KIF1A-SW and found to be similar (Supplementary Information; Fig. S7). This similarity indicates that the lower mean termination force is a consequence of the shorter engagement duration. Taken together, when loop-12 was substituted, KIF1A disengaged from the strong-binding state more readily under load. Interestingly, when subtilisin-treated microtubules were used in the three-bead assay to determine whether removal of the highly negatively charged C-terminal tail of tubulin (E-hook) produced a similar effect, we found that there was a large variability in the attachment duration for different microtubule dumbbells (Fig. S8). Although it is unclear whether this variability is due to absence of the E-hooks, nonspecific cleavage of other regions of tubulin, or some other effect, we would like to draw caution to the use of subtilisin-treated microtubules, especially in loaded assays. Comparison with recombinant tubulin lacking C-terminal tails should elucidate this aspect in the future.

**Figure 5:**
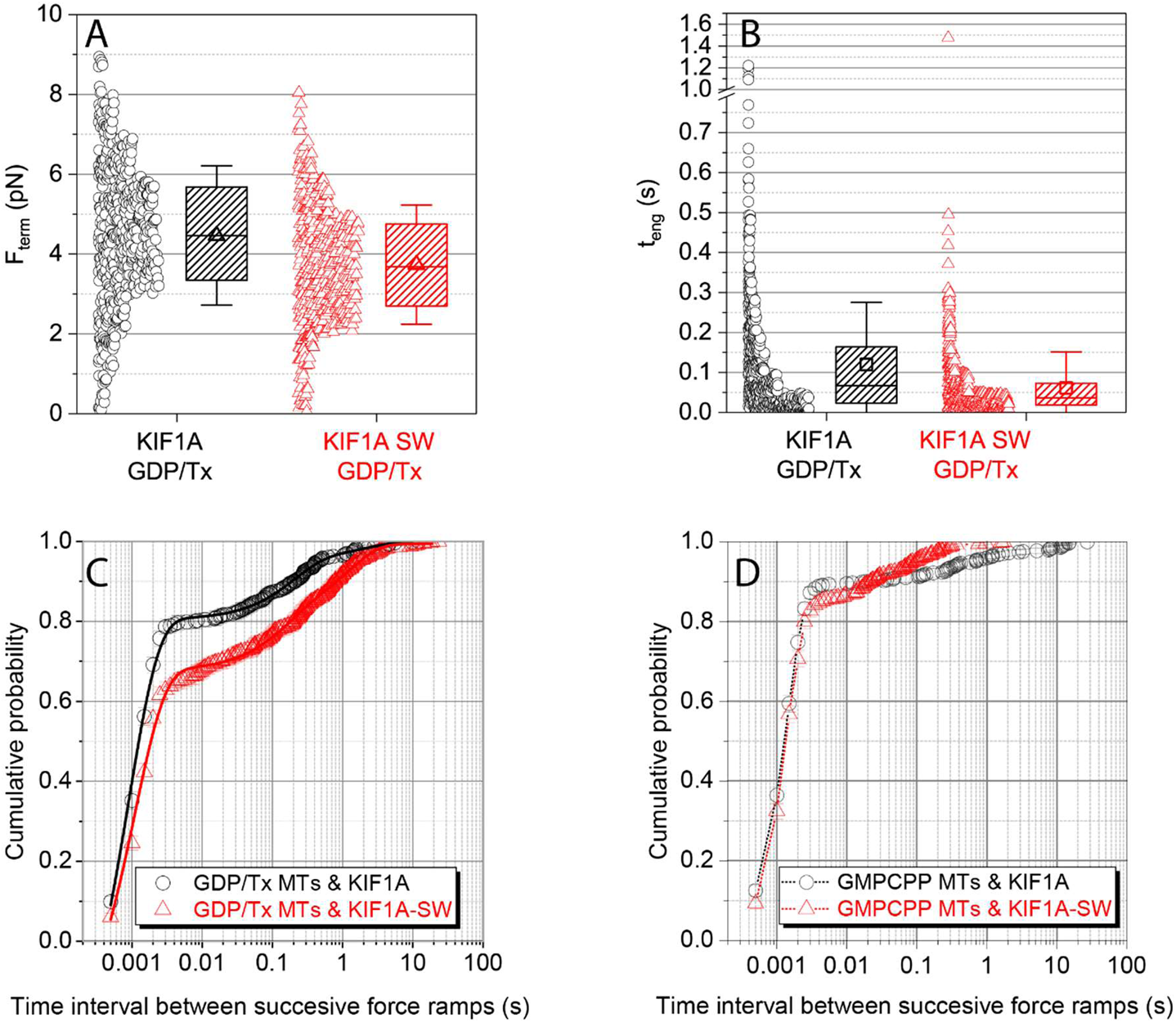
Influence of loop-12 and microtubule lattice on KIF1A performance under load. A) Comparison of KIF1A and KIF1A-SW termination forces, F_term_ for primary events on taxol-stabilized microtubules. (B) Comparison of KIF1A and KIF1A-SW engagement times, t_teng_ for primary events on taxol-stabilized microtubules. (C and D): Cumulative probability distribution of time intervals between successive force ramps, t_restart_ for KIF1A (open circles) and KIF1A-SW (open triangles) on (C) taxol-stabilized and (D) GMPCCP-stabilized microtubules. Data include both primary and secondary events. Error bars are calculated using the bootstrap method (27), the solid lines in (C) represent fitting to three-exponential decay function (see Materials and Methods; Table I) and the dotted lines in (D) are just linear connections between the data points to serve as guide to the eye.

To investigate whether loop-12 of KIF1A contributes to the motor’s ability to rapidly reengage following termination of a force ramp, we quantified the time before reengagement, t_restart_ for the loop swap mutant. We found that 79% of KIF1A events reengaged within 2 ms of disengagement, compared with only 66% of KIF1A-SW events (Fig. 5C). Coupled with the lack of effect on the unloaded landing rate (Fig. 4 E and F), positive charge in loop-12 does not significantly contribute to the initial interaction of the motor with the microtubule from solution, and it plays only a minor role in the fast reengagement with the microtubule following termination of a force ramp (Fig. 5C).

Our final investigation into the mechanism of fast reengagement kinetics of KIF1A asked whether properties of the microtubule lattice affect the KIF1A reengagement kinetics. Thus, instead of using taxol-stabilized GDP microtubules in the three-bead assay, we used microtubules polymerized with GMPCPP which have an expanded microtubule lattice (28–31). The distributions of termination forces and engagement times were not substantially impacted for either KIF1A or KIF1A-SW on GMPCPP microtubules (Fig. 5A and B; Table S2), suggesting that the dissociation rate of KIF1A under load is not affected by the nucleotide state of the microtubule. However, the probability of restarting within 2 ms increased on GMPCPP microtubules relative to Taxol/GDP microtubules (Fig. 5C and D). Strikingly, differences between the KIF1A and KIF1A-SW restart times that were observed on Taxol/GDP microtubules were abolished on GMPCPP microtubules. Shorter restart times on GMPCPP microtubules were also observed for KIF5B (Fig. S9). Thus, the rate of reengagement with the microtubule under load is affected by i) the identity of the motor, ii) the presence of the loop-12, and iii) the nucleotide state of the microtubule lattice.

## Discussion

The ability of kinesin motors to power intracellular transport against mechanical loads is integral to their function. The influence of load on motor speed and microtubule attachment lifetimes has been characterized using optical tweezers for a number of kinesin isoforms (e.g., (17, 32–34)). However, little is known about the load-dependence of kinesin-3 motility, which is of particular interest given its superprocessive behavior under zero load. Here, we find that KIF1A processive runs are readily terminated under load, resulting in lower average termination forces as compared to KIF5B. However, this behavior is compensated for by a rapid reengagement of the motor and recovery of force which is particularly apparent in the three-bead assay. These rapid KIF1A reengagement kinetics, also observed in a recent single-bead trap study (13, 35), are consistent with the fast bimolecular association rate constant for microtubule binding reported in a recent biochemical study (14). KIF1A therefore represents a different paradigm than KIF5B for an efficient transporter under force by rapidly and repeatedly reengaging with the microtubule and restarting its processive motion. Thus, whereas KIF1A is superprocessive in the absence of load, under load it may be better characterized as super-engaging.

### Performance of KIF1A under load

By implementing the three-bead assay in a dual-beam optical tweezers setup, we were able to investigate the performance of KIF1A as it stepped against loads oriented primarily parallel to the microtubule long axis. Importantly, we found that that KIF1A forces, although somewhat smaller on average, are comparable to those generated by KIF5B. KIF1A did not generate long-lived (> 0.2 s) force plateaus, or “stalls” seen frequently with KIF5B in the three-bead assay (19); instead, KIF1A more often disengaged before reaching a plateau. Thus, instead of quantifying a “stall force”, we quantified the termination force, F_term_, of the force ramps and found that in both the single- and three-bead assays, <F_term_> was smaller for KIF1A than for KIF5B. The lower KIF1A termination forces reflect the inability of KIF1A to remain strongly engaged with the microtubule under load, which may be a useful adaptation to achieve bidirectional motion (discussed below). Interestingly, the engagement times and termination forces for both KIF1A and KIF5B are smaller in the single-bead rather than in the three-bead assay, which demonstrates that vertical loads accelerate disengagement of these kinesin isoforms from the microtubule.

### Mechanism of fast KIF1A reengagement

A distinct feature of KIF1A motor behavior is its fast reengagement with the microtubule following the termination of a force ramp. Almost 79% of reengagements for KIF1A in the three-bead assay occurred within 2 ms whereas only 20% for KIF5B (Fig. 3B). Consensus models for the kinesin chemomechanical cycle point to the motor being in a weak-binding ADP-P_i_ or ADP state at the termination of the force ramp, and the transition to the strong-binding state to start the next force ramp requiring ADP release to generate the tight-binding apo state (36). Furthermore, two recent kinesin-1 optical trapping studies characterized the fast unbinding and rebinding events that occur while kinesin-1 slips backwards after it approaches its stall force (24, 25). Toleikis et al. found that during stall plateaus the bead slipped backward in 8 nm and longer displacements (24). Dwell times preceding backward displacements were longer than those preceding forward steps, consistent with the motor releasing P_i_ and slipping backwards in the ADP state. Using a small, high refractive index bead, Sudhakar et al found that during the backslipping process, the bead paused transiently (~30 μs) at 8 nm increments, consistent with the motor interacting transiently with successive tubulin subunits as it slid backwards along the protofilament (25). Both studies concluded that under load, kinesin-1 can enter a weakly-bound ADP or ADP-P_i_ state and slip backwards along the microtubule, and then reengage and recover. The drag coefficient of the microtubule dumbbell in our three-bead assay masks detection of microsecond interactions between KIF1A and the microtubule during the backward displacements. However, the millisecond-scale rescue of processive motion that we observe is consistent with KIF1A entering a weak-binding slip state like KIF5B, but transitioning back to a strong-binding, force-generating state much faster than KIF5B.

To explore the kinetics of this reengagement process, we constructed a kinetic model and fit it to our normalized cumulative distributions of restart times for KIF1A, KIF1A-SW, and KIF5B. In the model, the motor starts in a weakly-bound *Disengaged* state. The motor can then either transition to an *Engaged* state and continue to step against the load, or it can dissociate and enter a *Detached* state. Our experimental t_restart_ times correspond to the time it takes to transition from the *Disengaged* state to the *Engaged* state. To account for the two slower time constants in the t_restart_ distributions, we included two *Detached* states, with the idea that transitions into and out of these detached states may be influenced by the bead geometry and other experimental uncertainties. Note that in the model, the rate of the fast reengagement population as well as the relative proportion of fast reengagement events are determined by a kinetic race between the engagement rate constant, k_e_, and the two detachment rate constants, k_+d1_ and k_+d2_.

When we fit the model to the experimental data, the KIF5B reengagement rate k_e_ was 100 s^−1^, whereas the KIF1A reengagement rate was 990 s^−1^. Transition into the strongly-bound state is thought to be limited by ADP release (36). Published values for the ADP release rate of KIF5B in the absence of external loads range from 110 s^−1^ to 306 s^−1^ (37–40), which is close to the estimated k_e_ from our model (Fig. 6). However, the estimated value of k_e_ for KIF1A is more than twice the reported rate of ~350 s^−1^ for the ADP release when KIF1A is bound to the microtubule in a one-headed state in the absence of external load (14). How can we account for this fast KIF1A reengagement rate? One possibility is that this transition is load dependent, such that rearward load on the motor when it engages with the microtubule accelerates ADP release and thus the transition to a strong-binding state. Another consideration is that in the three-bead assay, the microtubule is under tensile forces even in the absence of interactions with kinesin; these tensile forces could alter the microtubule lattice in a way that enhances KIF1A engagement kinetics and/or ADP release. A third possibility is that KIF1A disengages in a nucleotide-free strong-binding state and is able to rapidly reengage without needing to release ADP or undergo the subsequent weak-to-strong transition. Additional experiments will be required to distinguish among these possibilities.

**Figure 6:**
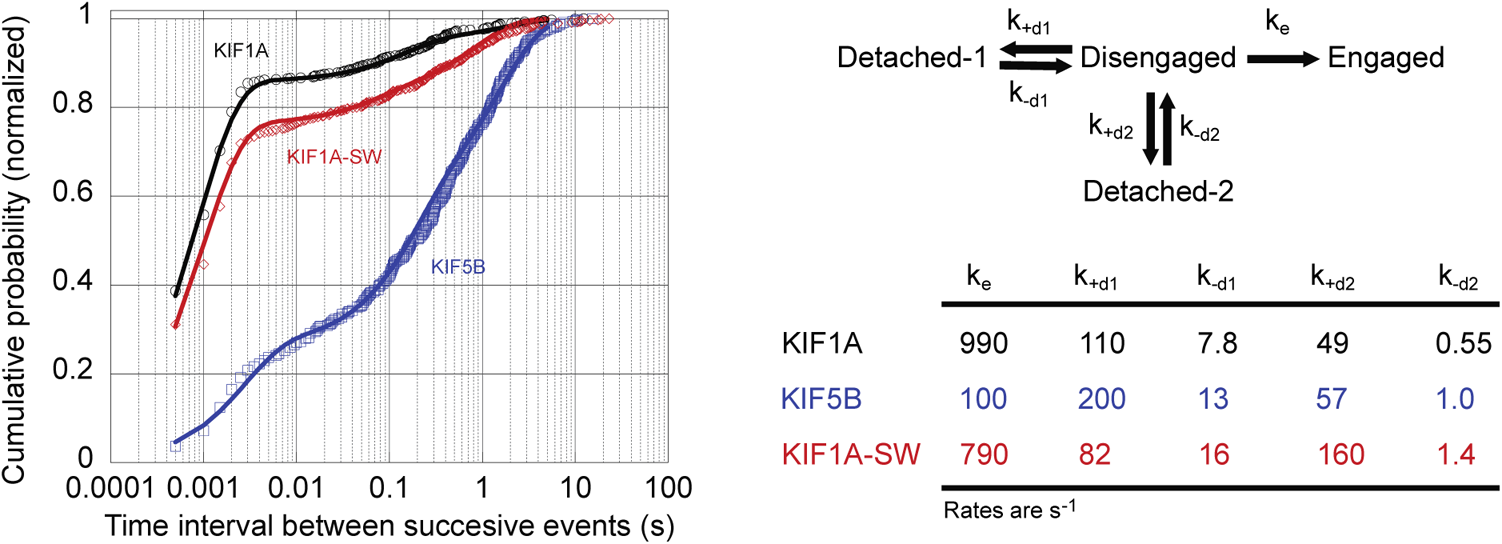
Comparison of reengagement rates between KIF1A and KIF5B. (A): Cumulative probability distribution of time intervals between successive force ramps, t_restart_ for KIF1A (red open circles), KIF1A-SW (blue open triangles), and KIF5B (green open squares) on taxol-stabilized microtubules. Data are offset to account for missed events resulting from the 0.5 ms minimum detection limit. Solid lines are fits to model of reengagement kinetics. (B): Kinetic model of motor reengagement. Following termination of a force ramp, the motor is in a Disengaged state. The motor can then reengage with the microtubule, with rate k_e_, or it can detach from the microtubule with two different rates k_+d1_ and k_+d2_ that depend on the motor-microtubule geometry and other factors. From the detached state, the motor can return to the disengaged state with rates k_-d1_ and k_-d2_ and reengage with the microtubule. Table shows the fits of the kinetic model to the t_restart_ times for the three motors.

### The role of the loop-12 in KIF1A motility

A distinctive feature of the KIF1A sequence is the positively charged K-loop insert of loop-12, but the role of this region in KIF1A motility under load remains murky. The importance of electrostatic interactions mediated by this loop was established in work on recombinant KIF1A monomers where it was shown that diffusive tethering by the K-loop enabled processivity (5, 6). In later work, KIF1A dimers were shown to be superprocessive at low ionic strengths (12 mM PIPES) (7, 8). As part of this work, it was shown that replacing the lysines in loop-12 of KIF1A with the analogous sequence from kinesin-1 (KIF5C) did not abolish the superprocessivity at low ionic strength, but it decreased the microtubule landing rate. We find here that in 80 mM PIPES buffer, which approaches physiological ionic strength, substituting the KIF1A loop-12 with that of *Drosophila* kinesin-1 decreases the unloaded run length six-fold, and deleting the C-terminal tail of tubulin has a similar effect. We propose that previous work (8) did not observe a change in run length due to the use of a very low ionic strength buffer. Consistent with this result, we found that KIF1A and KIF1A-SW had similar run lengths to each other in 12 mM PIPES buffer (Fig. S6).

We found that swapping the kinesin-1 loop-12 into KIF1A had only minor effects on the ability of KIF1A to remain engaged with the microtubule under load. For instance, in the three-bead assay, swapping loop-12 reduced F_term_ by ~25% and t_eng_ by ~2-fold. Importantly, the loop-12 swap only marginally affected the initial landing rate of the motor on the microtubule at the near-physiological ionic strength (80 mM PIPES) in the absence of load, as assessed by the single-molecule landing rate in TIRF and the bimolecular on-rate measured by stopped flow. Thus, when a motor in solution first encounters a microtubule, transition to the strong-binding state and ADP release is mediated by interaction of the canonical microtubule binding site (41) with the microtubule, rather than through initial formation of a tethered intermediate that is stabilized by the K-loop.

In the three-bead assay the fraction of reengagement events that occurred within 2 ms decreased by only ~25% for KIF1A-SW. This result is broadly consistent with the lack of an effect of the K-loop on the landing rate from solution. Intriguingly, when we quantified reengagement kinetics on microtubules polymerized in GMPCPP, which have been shown to have expanded lattices compared to Taxol/GDP microtubules (28, 30, 31), the proportion of rapid reengagement events increased for KIF1A-SW and matched that of wild-type. This result suggests that interaction of KIF1A with the microtubule under load are affected, albeit minimally, by both changes in charge in loop-12, as well as differences in the microtubule lattice. A recent study examining delivery of vesicles to synaptic boutons found that KIF1A has a lower affinity for GMPCPP microtubules compared to GDP/taxol microtubules (42). That reduced affinity was not observed in our measurements, but there are a number of differences between the assays, most notably load and concentrations of motors (single molecule *vs* saturating).

### Optical Trapping Geometry

Recent experimental (19) and theoretical (18) studies revealed the impact of optical trapping geometry on the measured parameters of KIF5B motility. Importantly, KIF5B stall times have likely been underestimated in the literature due to the vertical force components inherent to the assay. Additionally, the accelerated detachments in the single-bead assay have masked microtubule-to-microtubule heterogeneity in stall times. Thus, it was important to evaluate the KIF1A motility with both the single- and three-bead optical trapping assays. Like KIF5B, we observed larger t_eng_ and F_term_ values for KIF1A using the three-bead geometry, but the microtubule-microtubule variability was not seen. Most striking was the > 2-fold increase in the fraction of the reengagement events observed in the three-bead assay. By using the three-bead assay, we revealed that ~80% of all KIF1A detachments are followed by reengagement within 2 ms, compared to ~34% in the single-bead assay. These results have important implications for understanding the unique mechanism by which KIF1A sustains motility in the presence of obstacles and resisting mechanical loads.

### Insights into the biological function of KIF1A

The principal role of KIF1A in cells is vesicle transport and, unlike KIF5A, which transports cargo exclusively in axons, KIF1A transports cargo in both axons and dendrites (43–45). Much of this transport is bidirectional (46), meaning that KIF1A must both navigate diverse microtubule substrates, but also transport cargo against hindering loads generated by dynein. Although KIF1A has been characterized as a superprocessive motor in the absence of load, it is clear from Budaitis et al. (13) and our work that mechanical load more easily ends these processive runs, compared to KIF5B.

Interestingly, KIF1A has evolved kinetic features that allow it to be super-engaging. First, KIF1A has a 10-fold faster bimolecular on-rate, compared to kinesin-1 in the absence of load (14, 47). This fast on-rate is not mediated by the highly charged K-loop, but rather by other structural and mechanochemical features of the catalytic domain (15). Second, KIF1A has a very high probability of entering a strongly-bound state capable of initiating processive motility within 2 ms of disengaging from the microtubule under load. During intracellular transport, these features confer a distinct advantage because they increase the probability a motor will rebind to the microtubule to reinitiate transport following disengagement. This reengagement may allow for more robust transport because motors that disengage will rapidly resume motion along the original or a neighboring microtubule, testing for the best path to achieve movement. These adaptations of KIF1A mechanochemistry facilitate bidirectional transport and navigation around obstacles.

## Materials and Methods

### Protein Constructs and Purification

The KIF1A-WT construct (adapted from Addgene #61665 (10)) consists of the *R. norvegicus* KIF1A residues 1-393, followed by a GCN4 leucine zipper for dimerization and an eGFP tag. The KIF1A-SW was modified by swapping the native loop-12 (residues 288 – 308) of the KIF1A construct with the *D. melanogaster* KHC loop-12 sequence (GNKTHIPYRD). Both constructs have a C-terminal His tag and were bacterially expressed and purified by nickel gravity column chromatography, as described previously (14). The elution buffer, consisting of 20 mM phosphate buffer, 500 mM sodium chloride, 500 mM imidazole,10 μM ATP and 5 mM DTT was supplemented with 10% glycerol before flash freezing and storing at −80 °C. Concentrations were determined using GFP absorbance at 488 nm.

Unlabeled porcine tubulin and its labeled analogues, (TRITC and biotin), GTP and Paclitaxel were purchased from Cytoskeleton, Inc. Mouse monoclonal anti-6xHis tag antibody and rat tubulin antibody which recognizes the C-terminal tail of α tubulin were purchased by ABCAM. GMPCPP was purchased from Jena Biosciences, Germany. Streptavidin coated polystyrene beads 1% w/v (0.82 μm in diameter) and silica microspheres 9,92 % solid w/v (5.0 μm in diameter) were purchased from Spherotech, (Lake Forest, IL). Amyl acetate and 2% Colloidon in amyl acetate were purchased from Electron Microscopy Sciences, PA. Glass coverslips 22 x 45 x 1.5 mm were purchased from Fisher Scientific. Glucose oxidase from *Aspergillous niger*, aqueous solution of catalase from bovine liver, dimethyl sulfoxide (DMSO), phenylmethylsulfonyl fluoride (PMSF), ATP, MgCl_2_ and Subtilisin A *Bacillus licheniformis* were purchased from Sigma Aldrich. Mouse anti-tubulin b3 antibody which recognizes the C-terminal tail of β tubulin was purchased from Bio-Rad Laboratories.

### Optical Tweezer Experiments

Taxol-stabilized GDP microtubules and GMPCPP-stabilized microtubules were prepared from non-polymerized porcine tubulin as previously described (19). For the single-bead assay, 4% TRITC-tubulin was included, while for the three-bead assay 4% TRITC tubulin as well as 48% biotinylated tubulin were included.

For the single-bead assay, nitrocellulose-coated coverslips were assembled into flow chambers of 20 μL volume as described previously (19), and used within 24 h of preparation. Aqueous solutions in BRB80 pH 6.9 were introduced in the chamber in the following sequence: 20 μL of 0.05 mg/mL anti-tubulin antibody (Bio-Rad Laboratories) for 5 min, 50 μL of 2 mg/mL casein for 4 min, 4 x 25 μL of 125 nM 4% TRITC microtubules supplemented with 2 mg/mL casein and 20 mM taxol for 4 x 1 min, wash with 100 μL of 2 mg/mL casein, and 50 μL of final solution containing kinesin beads, 2 mM ATP, 2 mM MgCl_2_, 50 mM DTT, 20 μM taxol, 5 mg/mL glucose, 1500 units/mL glucose oxidase, and 0.2 units/mL catalase. The open ends of the flow chamber were sealed with vacuum grease to prevent evaporation during the experiment. To ensure single-molecule interactions concentrations of kinesin were used such that no more than one out of three kinesin-decorated beads interacted with surface immobilized microtubules.

For the three-bead assay, a solution of silica spherical pedestals (dia. 5.0 μm) was dried on a coverslip, coated with nitrocellulose-film, and assembled into ~ 20 μl flow chambers, as previously described (19). Aqueous solutions in BRB80 were introduced into the flow chamber in the following sequence: 20 μL of 0.2 mg/mL anti-6xHis antibody (Abcam) for 5 min, 50 μL of 2 mg/mL casein for 4 min, 50 μL of kinesin construct ~1 nM supplemented with 2 mg/mL casein for 5 min, 100 μL of 2 mg/mL casein wash, and 50 μL of final solution containing 5 nM 48% biotinylated-4% TRITC microtubules, 2 mM ATP, 2 mM MgCl_2_, 50 mM DTT, 20 μM taxol (excluded when GMPCPP microtubules were used), 5 mg/mL glucose, 1500 units/mL glucose oxidase, and 0.2 units/mL catalase. Before sealing the chamber with vacuum grease, 3–4 μL of streptavidin beads (dia. 0.82 mm) diluted 1:30 in final solution without microtubules were introduced from one side of the chamber. To ensure single-molecule interactions concentrations of kinesin were used such that no more than one out of three kinesin-decorated spherical immobilized pedestals interacted with microtubule dumbbells.

### Optical Tweezer Instrumentation and Data Analysis

We used a custom made a dual-laser beam (1064 nm) optical trap system equipped with a 63x water objective, 1.2 numerical aperture as previously described (19). The trap stiffness (pN/nm) and the system-calibration factor (pN/V) for each trapped bead were determined in the absence of any microtubule interaction by calculating and fitting to a Lorentzian function the power spectrum of the Brownian motion of the beads in the trap. Microtubule dumbbells were subjected to stretching forces of 4-5 pN by moving the two laser beams apart. The trap stiffness of the individual laser beams for single-bead assays was 0.04–0.12 pN/nm and for three-bead assays was 0.060–0.090 pN/nm. The higher total stiffness in the three-bead assay was required to accommodate the sum of the stretching forces on the microtubule dumbbell and the forces generated by kinesin. The higher stiffness also decreases the relaxation time of the dumbbell close to the relaxation time of the bead in the single-bead assay (see Supplementary Methods). Since the laser traps are stationary, a piezoelectric stage controller was used to move the flow chamber and therefore control the relative position between single beads and surface-immobilized microtubules or between microtubule dumbbells and surface immobilized spherical pedestals. Data were digitized at a scanning rate of 2 kHz and filtered at 1 kHz using in-house software written in LabVIEW. Strictly monotonic decrease in the force trace were considered as disengagement events when the size of decrease in force was higher than the standard deviation of a 3 ms window either right before or right after the monotonic decrease event.

For data analysis, in-house software written in LabVIEW was used, while for statistical analysis, curve fitting and graphs Origin 2018b software was used, as described previously (19). The cumulative probability distributions for the time intervals between successive force ramps were fit using the tri-exponential decay function:

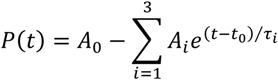

All the parameters were free, except *t*_0_ which was set equal to 0.5 ms and corresponds the temporal resolution of the optical tweezers’ data. The final amplitude values reported in Table I are relative values divided by their total sum ΣA_i_ such that the probability density is normalized to 1 over the observed range of values t ≥ t_0_ instead of t ≥ 0, and A_0_ = 1 (27, 48, 49). The kinetic modeling in Fig. 6 was done using the kinetics simulator Tenua (http://bililite.com/tenua/).

### TIRF Experiments

Single-molecule tracking of GFP-labeled KIF1A-WT and KIF1A-SW were performed on a Nikon TE2000 TIRF microscope at 21 °C, as described previously (40, 50, 51). Flow cells were prepared by flowing in 2 mg/ml casein, followed by full-length rigor kinesin (40) and taxol-stabilized, Cy5 (GE Healthcare) labeled microtubules. The microtubules were incubated for 30 sec, followed by a wash, and repeated 2x. Motors were diluted to 200-500 pM and added to the flow cell in the presence of 2 mM ATP and imaged at 5 fps. The kymographs were analyzed manually using Fiji (NIH) (52) to determine the run lengths, velocities and landing rates.

### Stopped Flow Experiments

Stopped-flow experiments were performed using an Applied Photophysics SX20 spectrofluorometer at 25 °C in BRB80 buffer, as previously described (14, 53). For k_on_^Mt^ measurements, a solution of 150 nM motor dimers and 0.25 mM free mADP was flushed against a solution containing 5 μM Taxol, 1 mM ATP, varying concentrations of taxol-stabilized microtubules (all final chamber concentrations). After mixing, mADP released from the bound head produced a decrease in fluorescence at 356 nm, which was fit with a single exponential to determine the k_obs_ at each microtubule concentration. The averaged trace of 5–7 consecutive shots was fit and reported for each trial. Linear fit to the rates versus the microtubule concentration gives the bimolecular on-rate (14, 53).

## Acknowledgements

W.O.H. thanks the University of Pennsylvania Physiology Department for hosting a sabbatical visit in Fall, 2019. This work was supported by NIH grant 5RM1GM136511 and National Science Foundation Science and Technology Center CMMI: 15-48571 to E.M.O., and NIH grants R01GM076476 and R35GM139568 to W.O.H, and F32GM137487 to A.M.G.

## Supplementary Information

### Supplementary Methods

#### Theoretical estimation of drag coefficients and relaxation times

The drag coefficient γ_microsphere_ of a microsphere with radius r_microsphere_ = 410 nm in an aqueous solution (viscosity coefficient η = 10^−9^ pN·s/nm^2^) is given by the Stokes’ law (1) as γ_microsphere_ = 6πηr _microsphere_ = 0.77 × 10^−5^ pN·s/nm. The drag coefficient γ_dumbbell_ of the microtubule dumbbell can be considered to a first approximation as the sum of the drag coefficients of the two beads (r = 410 nm) and that of a microtubule segment of length L = 10 μm (2). Approximating the microtubule as a solid cylinder of radius R = 12.5 nm, the drag coefficient for motion parallel to the cylindrical axis is γ_microtubule_ = 2πηL/[ln(L/(2R))-0.2] (1), and therefore γ_dumbbell_ = 2·γ_microsphere_ + γ_microtubule_ = 2.7 × 10^−5^ pN·s/nm = 3.5·γ_microsphere_. Since the characteristic relaxation time of a laser trapped object is τ = γ/k, where k is the stiffness of the laser trap, the microtubule dumbbell will have similar τ to that of a microsphere only if trap stiffness k~0.04 pN/nm is increased by the same factor as the drag coefficient, i.e 3.5 x 0.04 pN/nm = 0.14 pN/nm.

#### Calculation speed as a function of force

To calculate the profile of speed as a function of force two different but equivalent approaches were used:

1. The average of the displacement ramps as a function of time was calculated as previously described (3), and was smoothed using a Savitzky-Golay filter with a 20-point window. The speed was calculated from the first derivative of the smoothed average displacement trace. The corresponding force value was calculated by multiplication of the average displacement value with the optical trap stiffness.
2. The speed was calculated for each displacement ramp by linear fit of successive 10 ms window segments. The corresponding force value was calculated by the average displacement value of each segment multiplied by the optical trap stiffness.

Both methods gave similar results as can be seen is Fig. S6.

#### Subtilisin treatment of polymerized microtubules

We adopted a modified version of the protocol from Rodionov et al. (4). Taxol-stabilized GDP microtubules (50 μM tubulin) were incubated with 1 μM of A subtilisin for 4 hours at 37 °C water bath. The reaction was then blocked by the addition of PMSF to final concentration of 2 mM and the sample was kept at RT for at least 1 hour before using it for further experiments. We confirmed by western blot that for shorter incubation periods the C-terminal tail of α-tubulin is not fully cleaved as has been reported previously (5) (Fig. S5). Primary antibodies specific for the C-terminal tail of α and β tubulin were used (see Materials and Methods).

### Supplementary Tables

**Table S1.**

| Assay | Construct | % Secondary Events |  |
| --- | --- | --- | --- |
|  |  | Taxol/GDP MTs | GMPCPP MTs |
| Single-bead | KIF1A | 11 | - |
|  | KIF5B | 10 | - |
| Three-bead | KIF1A | 39 | 44 |
|  | KIF5B | 13 | 22 |
|  | KIF1A-SW | 26 | 49 |

**Table S2.**

| Assay | Construct | Primary Events<br>Taxol/GDP MTs |  | Secondary Events<br>Taxol/GDP MTs |  | Primary Events<br>GMPCPP MTs |  | Secondary Events<br>GMPCPP MTs |  |
| --- | --- | --- | --- | --- | --- | --- | --- | --- | --- |
| | | $\langle F_{\text{term}} \rangle$<br>(pN) | Med-$t_{\text{eng}}$<br>(s) | $\langle F_{\text{term}} \rangle$<br>(pN) | Med-$t_{\text{eng}}$<br>(s) | $\langle F_{\text{term}} \rangle$<br>(pN) | Med-$t_{\text{eng}}$<br>(s) | $\langle F_{\text{term}} \rangle$<br>(pN) | Med-$t_{\text{eng}}$<br>(s) |
| 1-bead | KIF1A | $2.2 \pm 0.99$ | 0.062 | $2.6 \pm 0.77$ | 0.023 | - | - | | |
| 3-bead | KIF1A | $4.1 \pm 1.8$ | 0.069 | $5.0 \pm 0.11$ | 0.067 | $3.2 \pm 1.3$ | 0.065 | $4.3 \pm 1.5$ | 0.079 |
| | KIF1A-SW | $3.5 \pm 1.4$ | 0.039 | $4.7 \pm 1.4$ | 0.030 | $3.4 \pm 1.5$ | 0.033 | $4.9 \pm 1.6$ | 0.043 |

**Table S3.**

| Construct | $\langle \text{Speed} \rangle$ ($\mu\text{m/s}$) | $\langle \text{Run Length} \rangle$ ($\mu\text{m}$) | Med-$t_{\text{eng}}$ (s) |
| --- | --- | --- | --- |
| KIF1A | $1.1 \pm 0.38$ | $6.3 \pm 4.2^{(*)}$ | 3.75 |
| KIF1A-SW | $1.3 \pm 0.42$ | $1.1 \pm 0.56$ | 0.73 |
| KIF1A & Subtilisin MTs | $1.3 \pm 0.39$ | $1.8 \pm 1.1$ | 1.3 |

(\*) The value has been corrected for events in which the motor runs all the way to the end of the microtubule (6).

### Supplementary Figures

**Figure S1:**
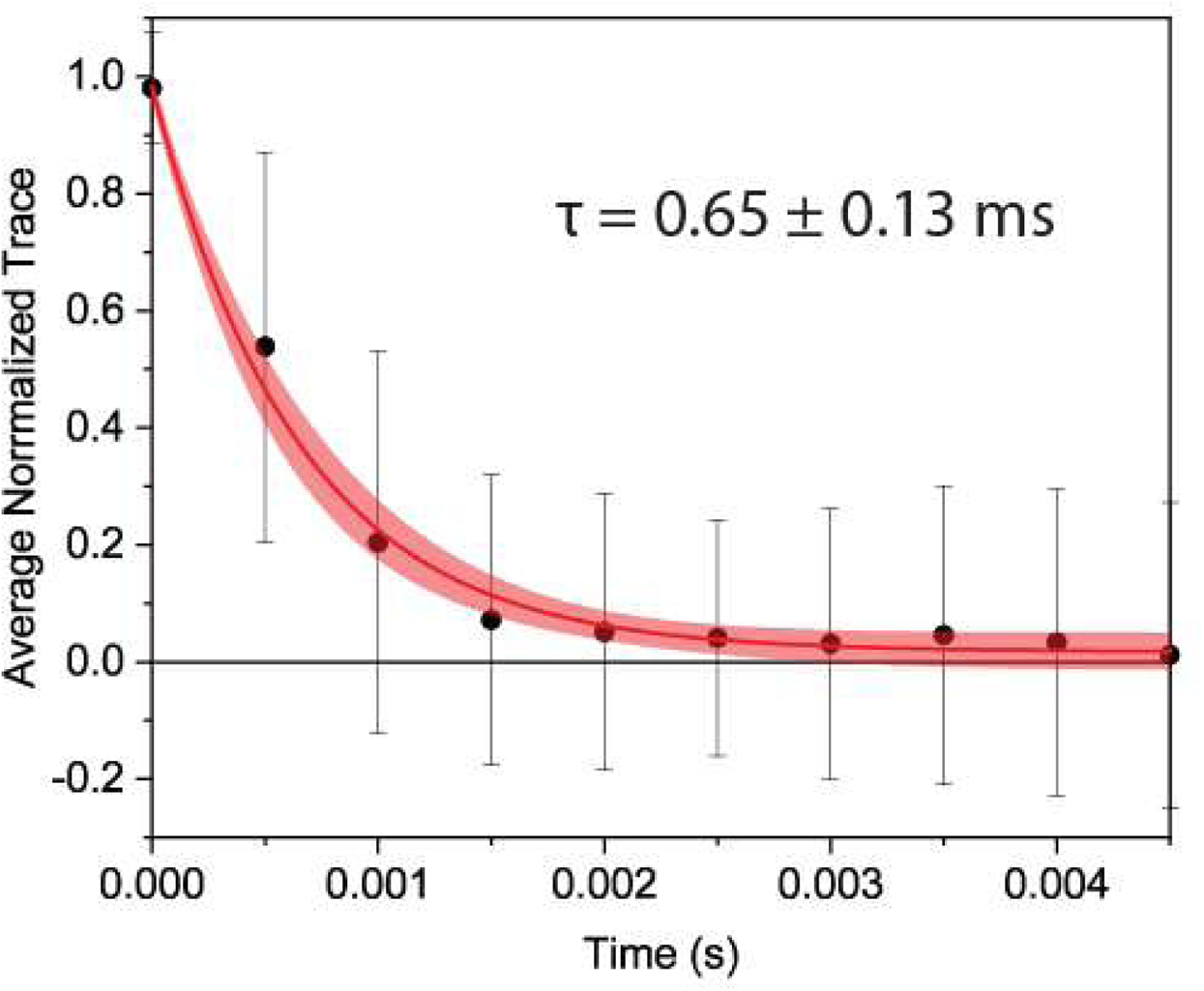
Average relaxation trace for 3-Bead assay. Relaxation traces from one dumbbell that correspond to either primary or secondary events and were strictly monotonic within 5 ms were normalized and then averaged (scatter points). Error bars correspond to standard deviations. The red line corresponds to single-exponential curve fitting and the 95% confidence band is indicated by the lighter red color.

**Figure S2:**
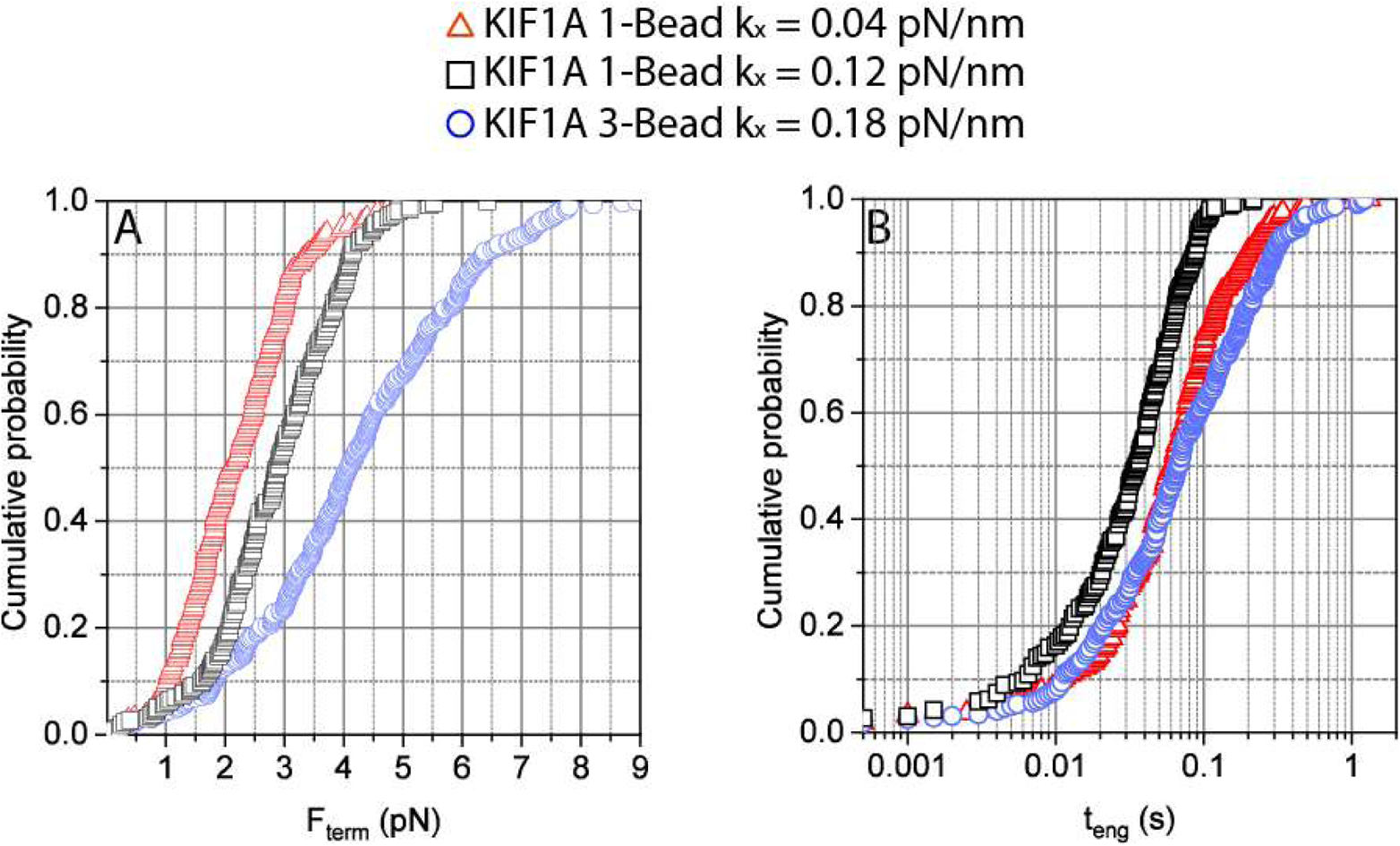
Comparison of 1-bead data at higher and lower stiffness, with the 3-bead assay. Cumulative probability distributions of terminal forces (A) and force ramp durations (B) for the single-bead and three-bead assays. To test whether larger the F_term_ in the three-bead assay (blue, open circles) was due to the higher trap stiffness, the trap stiffness was tripled in the single-bead assay from 0.04 pN/nm (red, open upright triangle) to 0.12 pN/nm (black, open square). Although there was a moderate shift toward the three-bead values, Fterm values are still clearly smaller in the one-bead assay. For t_eng_, increasing the single-bead trap stiffness closer to the three-bead value shortened t_eng_, reflecting the faster dissociation rates under load in the single-bead assay.

**Figure S3:**
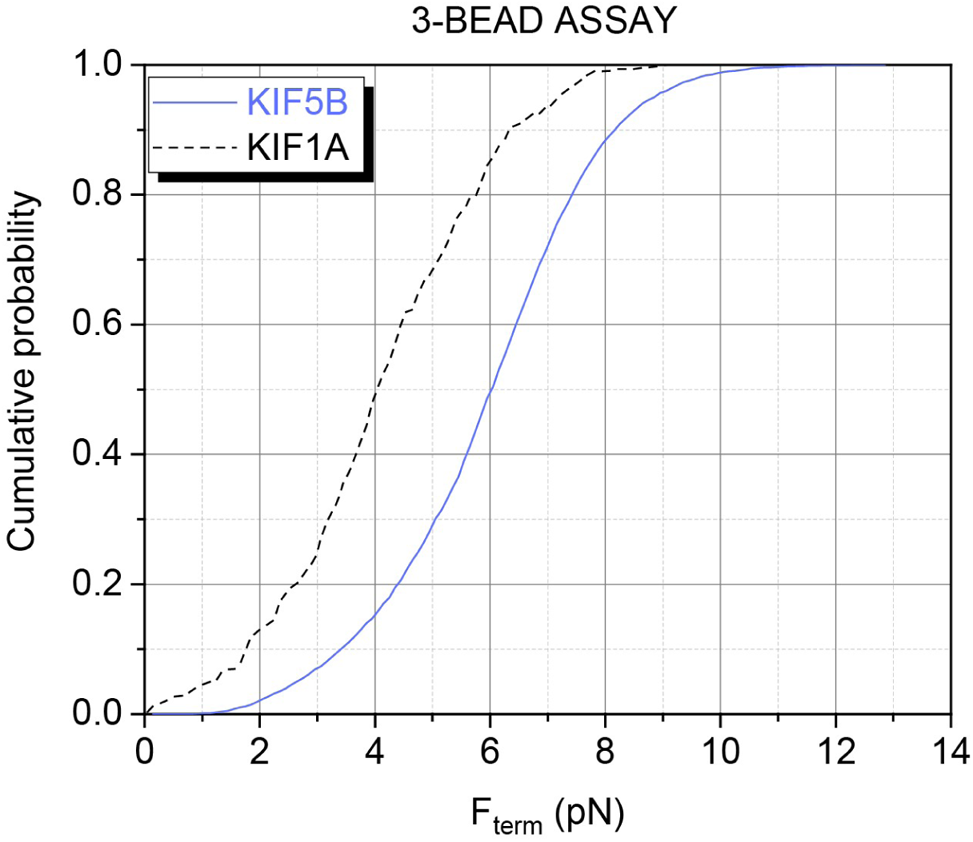
Cumulative probability of F_term_ for KIF1A and KIF5B in the three-bead assay.

**Figure S4:**
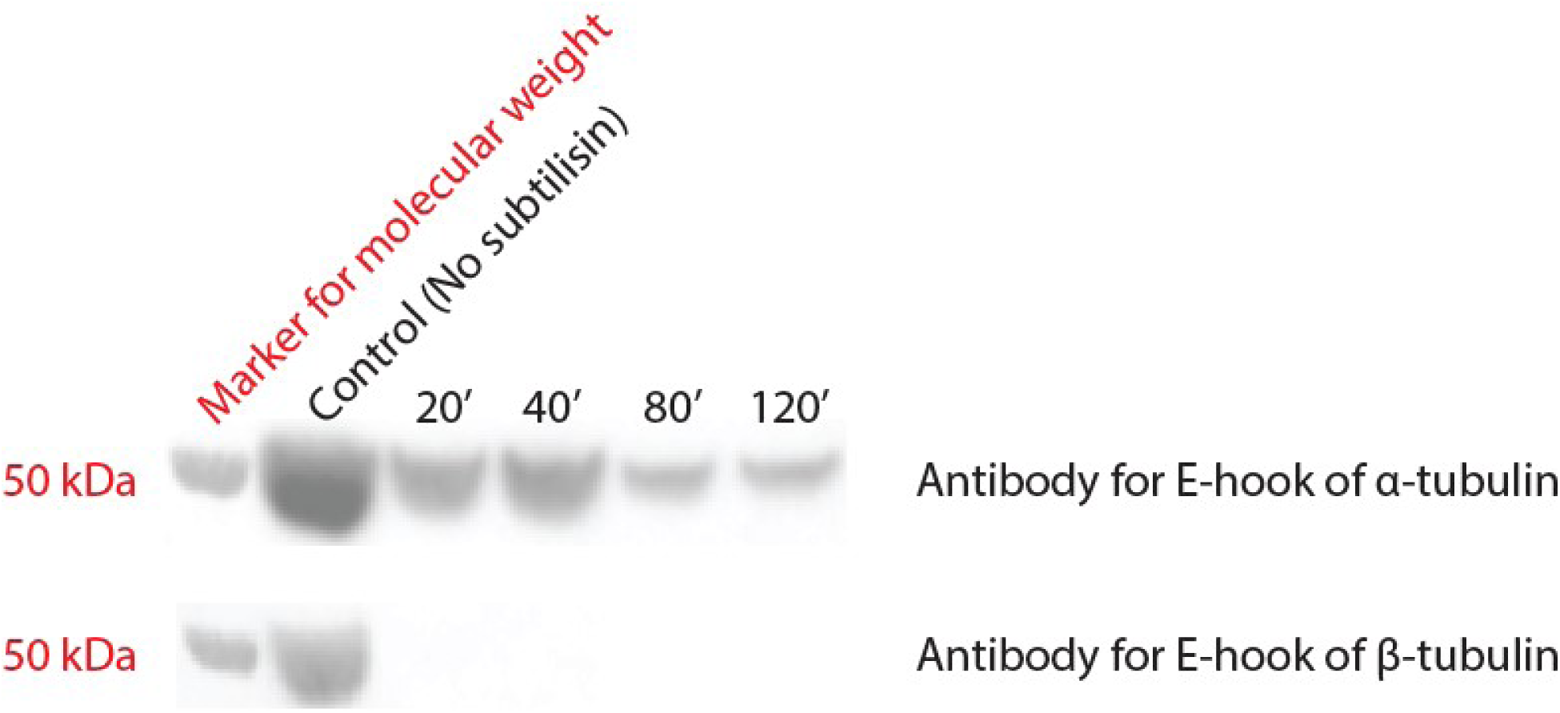
Time-course of subtilisin mediated cleavage of tubulin’s C-terminal tails from polymerized microtubules stabilized with Taxol. Western-blot against the C-termini of α and β tubulin after treatment of polymerized microtubules with subtilisin at 37 °C for 20, 40, 80 and 120 min (see Supplemental Materials and Methods).

**Figure S5:**
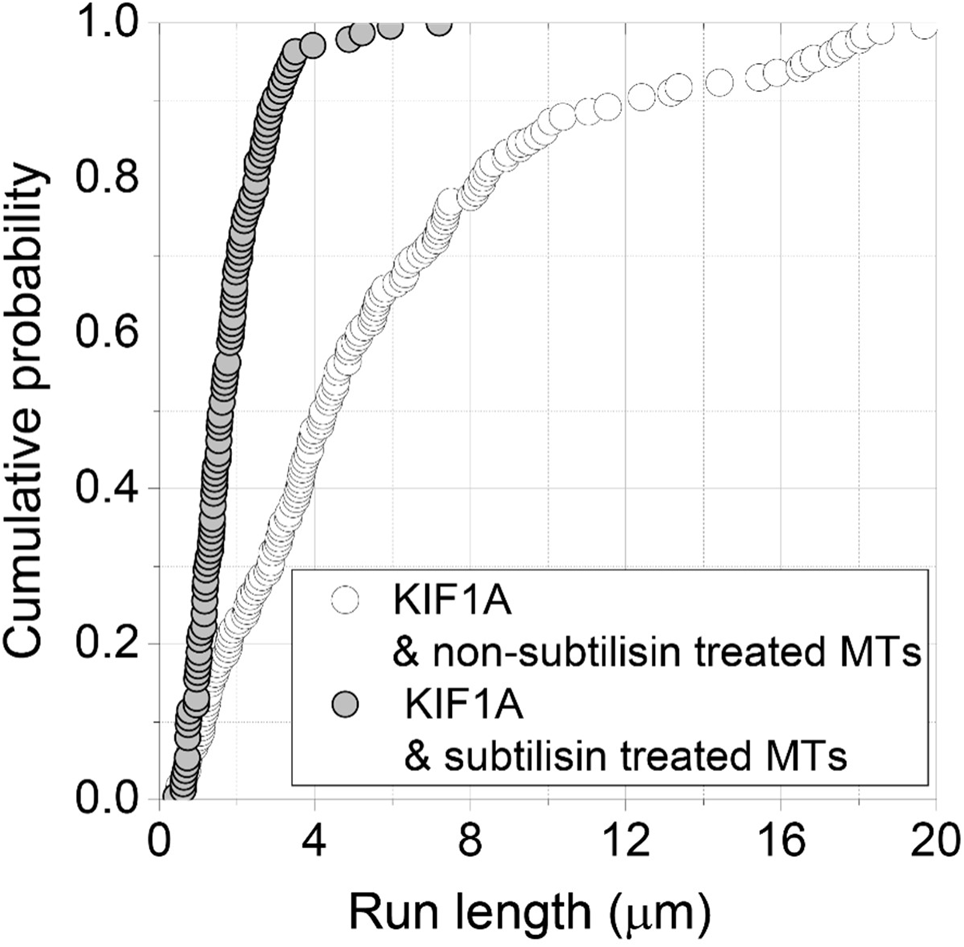
Unloaded run length of KIF1A on MTs that were treated and not with subtilisin. Cumulative probability of the run length for single KIF1A molecules observed under TIRF microscopy in BRB 80 buffer when microtubules were treated with subtilisin (gray filled circle) and not (open black circles).

**Figure S6:**
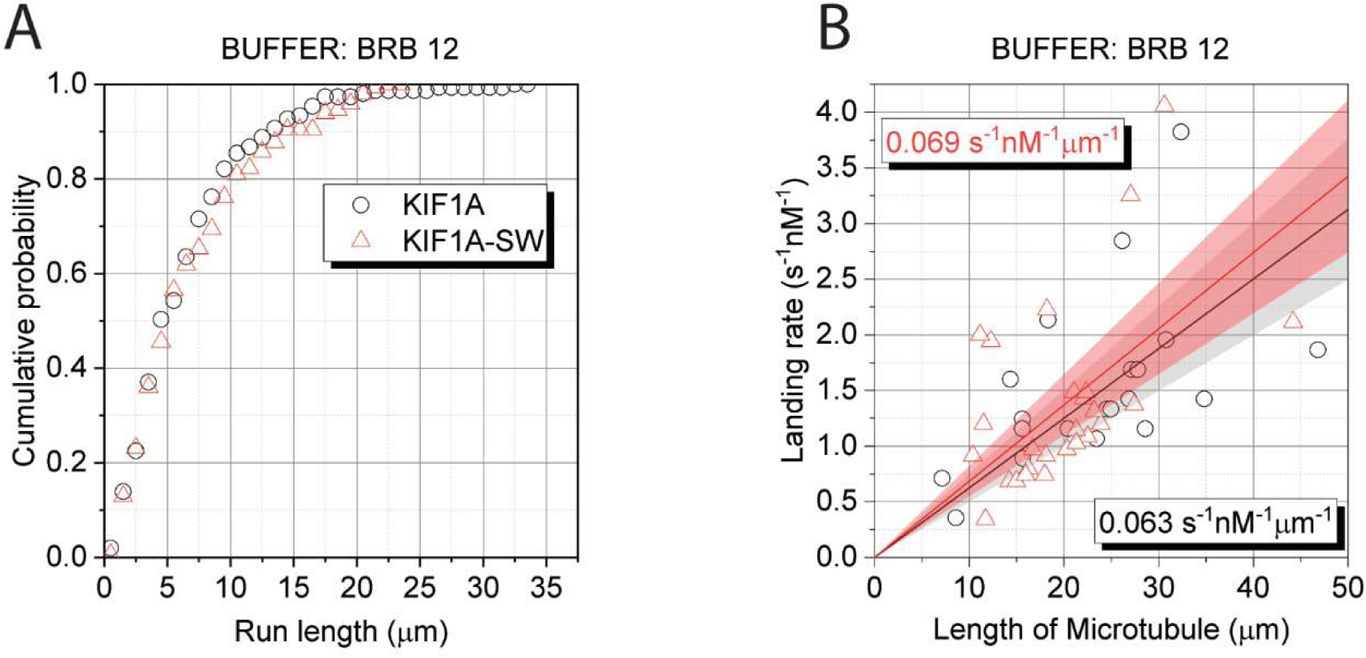
Run length and landing rate of KIF1A and KIF1A-SW in buffer BRB12. (A) Cumulative probability of the run length and (B) landing rates for single KIF1A (black) and KIF1A-SW (red) molecules observed under TIRF microscopy in BRB12 buffer.

**Figure S7:**
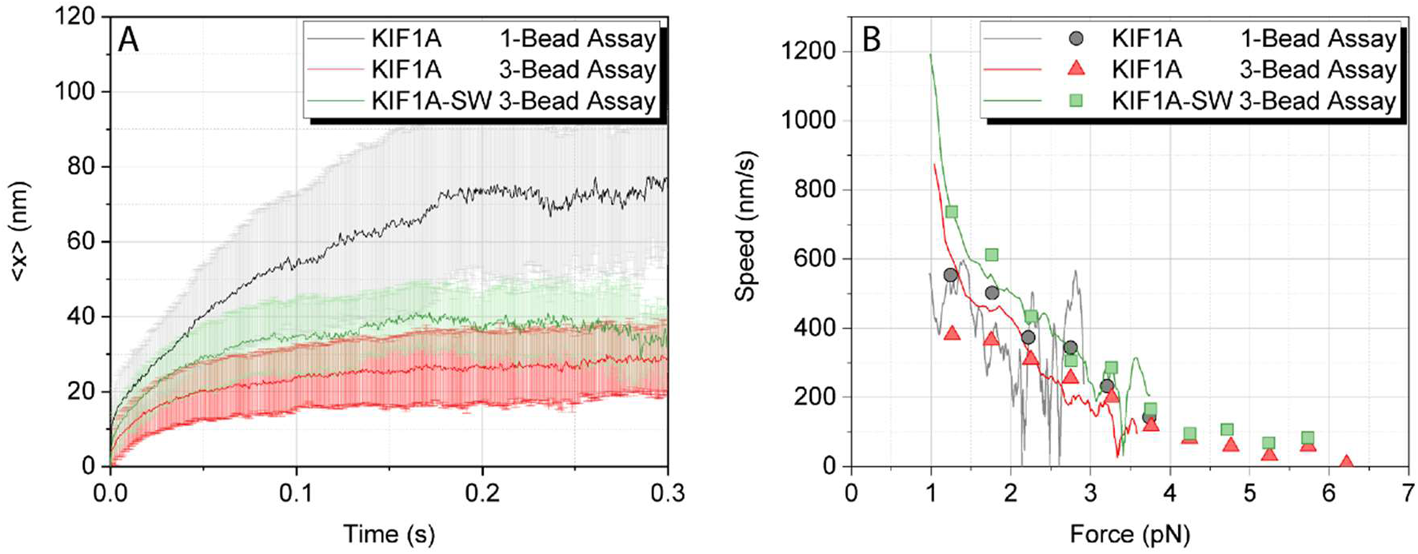
Ensemble trajectories and force-velocity curve. A. Ensemble average of displacement ramps for single- and three-bead assays and KIF1A constructs using only the primary events, B. Speed as a function of force calculated either from the derivative of the ensemble average trace (continuous line) or by piecewise calculation of the velocity for each displacement ramp over a 10 ms time window (not sliding) and subsequent averaging over all displacement ramps for each assay and KIF1A construct (scatter points), as described in Supplemental materials and methods.

**Figure S8:**
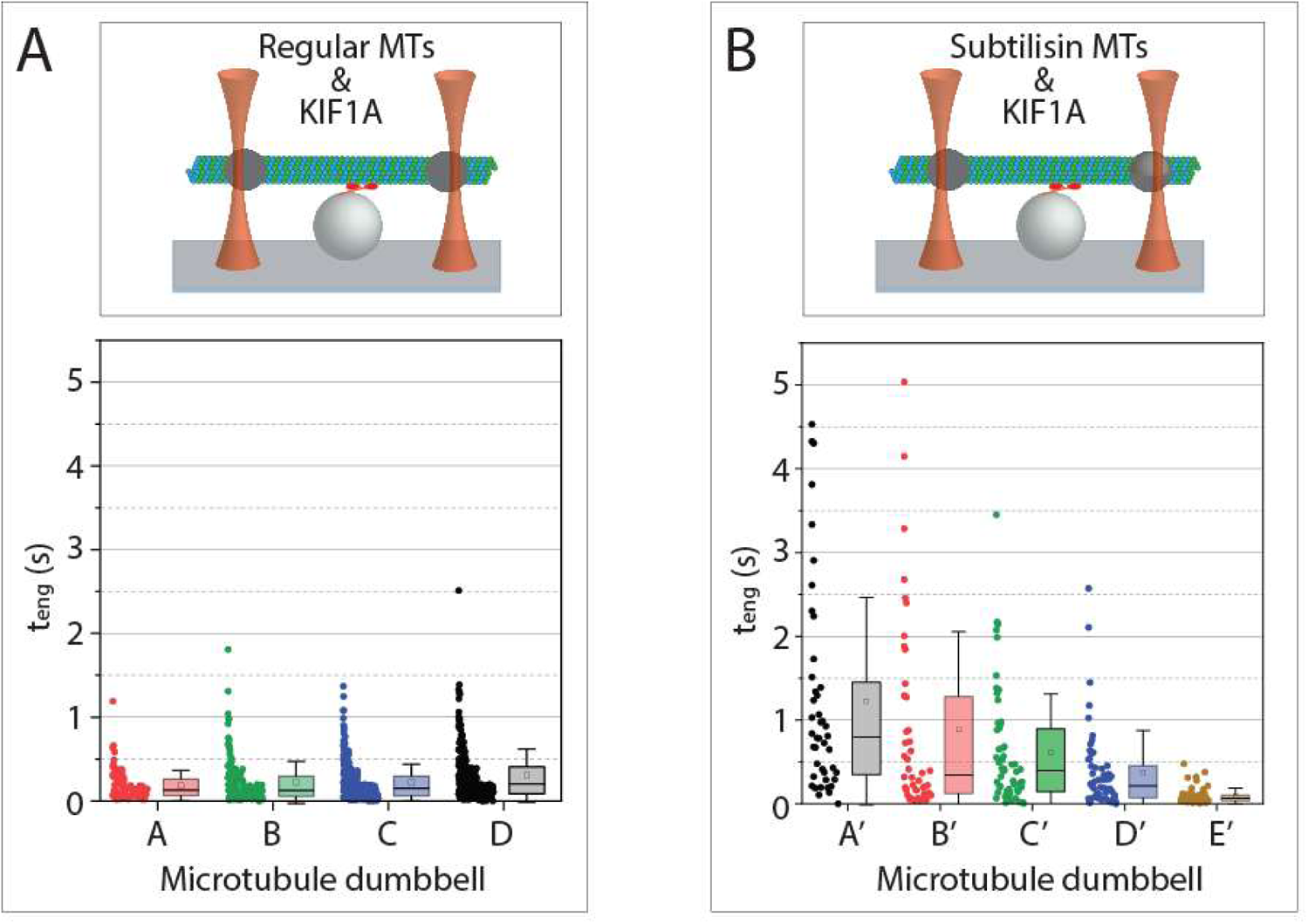
Three-bead assay using subtilisin treated microtubules. Statistics-box plot of the attachment durations t_eng_ between KIF1A single-molecules and (A) microtubule dumbbells not treated with subtilisin and (B) microtubule dumbbells after treatment with subtilisin A (see Supplemental Materials and Methods).

**Figure S9:**
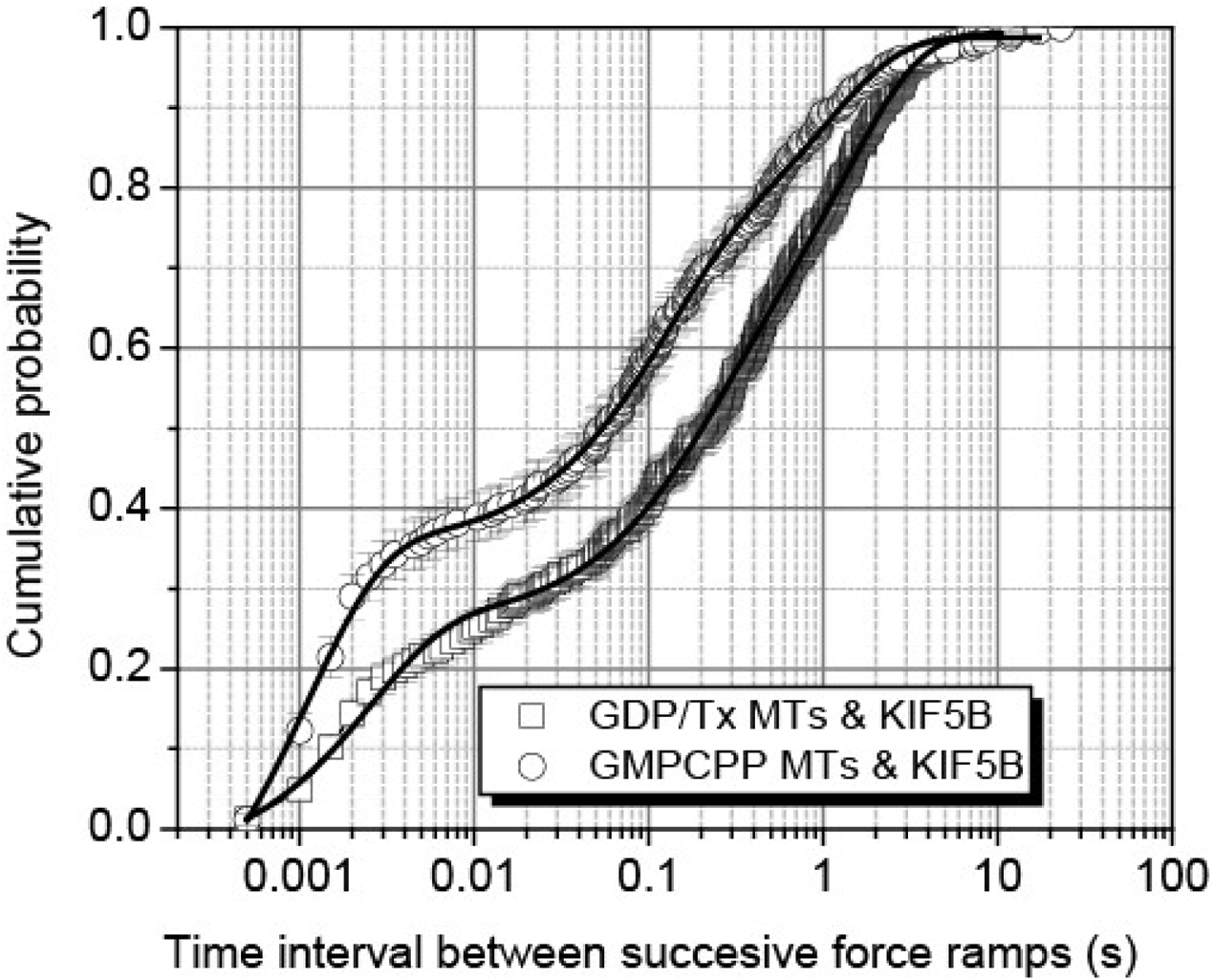
Comparison of KIF5B restart times on different microtubules. Cumulative probability plot of the time intervals between successive force ramps for K560 on taxol (open squares) and GMPCCP (open circles) stabilized microtubules, using the three-bead assay. Error bars are calculated using the bootstrap method (7) and the solid lines represent fitting to three-exponential decay function (see Materials and Methods).

